# EZH2 Inhibition Induces a Metabolic Stress Response Sensitizing TNBC to Glutaminase Targeting

**DOI:** 10.1101/2025.09.04.674332

**Authors:** Lucas Porras, Marina Carrière, Ann-Sophie Gironne, Elise Quadri, Gabriel Alzial, Hugo Philippeau, Yousef Aleassa, Annie Monast, Faustine Gorse, Myriame Saint-Arnaud, Mariana De Sa Tavares Russo, Sylvie Mader, Daina Avizonis, Sébastien Lemieux, Morag Park, Geneviève Deblois

**Author notes:** ^7^ Lead contact (Geneviève Deblois).

## Abstract

EZH2, the catalytic subunit of Polycomb Repressive Complex II, is overexpressed and associated with poor prognosis in triple-negative breast cancer (TNBC). Although EZH2 inhibition induces significant changes in chromatin landscapes and gene expression, it has a limited impact on the growth of TNBC models, suggesting adaptive compensatory mechanisms. Here, we demonstrate that EZH2 inhibition leads to the accumulation of misfolded proteins and double-stranded RNA (dsRNA), triggering an essential integrated stress response (ISR) through PKR and PERK activation. By inducing ISR-mediated ATF4 activation, EZH2 inhibition enhances amino acid flux and promotes glutaminolysis, in turn activating mTOR signaling to support TNBC cell survival. Pharmacological targeting of this metabolic adaptation with a glutaminase (GLS) inhibitor in combination with EZH2 inhibition significantly impairs TNBC cell proliferation and tumor growth. These findings reveal a stress-driven metabolic adaptation that sustains TNBC survival upon EZH2 blockade and highlight inhibition of this pathway as a strategy to enhance the efficacy of EZH2 inhibitors in TNBC.

## INTRODUCTION

Epigenetic reprogramming underlies key tumorigenic processes, including epithelial-to-mesenchymal transition (EMT)^1^, immune evasion^2^, metastasis^3^, and drug resistance^4–6^. Epigenetic regulators thus represent promising therapeutic targets to disrupt tumor progression. The Polycomb Repressive Complexes 1 and 2 (PRC1/PRC2) are key components of the Polycomb Group (PcG) proteins, which regulate chromatin accessibility by forming facultative heterochromatin domains^7^. PRC2 establishes transcriptionally repressive chromatin states through the methylation of histone H3 at lysine 27 (H3K27me1/2/3), a reaction catalyzed by its SET domain-containing methyltransferases Enhancer of Zeste Homolog 1 and 2 (EZH1/EZH2). PcG complexes are crucial for cell-fate determination during embryogenesis and differentiation^8^.

Dysregulation of PcG-mediated silencing, due to mutations or aberrant expression, is common in cancer^9,10^ and has been associated with tumorigenesis^11^. EZH2 is frequently overexpressed in solid tumors, including breast, ovarian, endometrial, prostate, liver, lung cancers, and melanoma, and is assocaited with metastasis and drug resistance^12,13^. Somatic mutations in EZH2 are predominantly observed in hematologic malignancies, with gain-of-function mutations driving oncogenic activity in lymphomas^14^ and loss-of-function mutations linked to tumor suppressor activity in T-cell leukemias and myelodysplastic syndromes^15,16^. Additionally, EZH2 activity is regulated by post-translational modifications, microRNAs, and cofactor interactions, some of which occur independently of PRC2, underscoring the complexity of EZH2 signaling in tumors. Although EZH2 represents an attractive anticancer target, in vivo preclinical studies have shown variable inhibitor activity and frequent resistance. More pronounced effects are observed in hematologic malignancies, particularly germinal-center lymphomas with EZH2 mutations, adult T-cell leukemia/lymphoma, and multiple myeloma^17–19^. By contrast, preclinical solid-tumor models show limited benefit from EZH2 inhibition, with clearer responses in specific genetic backgrounds^20,21^. The EZH2 inhibitor tazemetostat (TAZVERIK) was approved for epithelioid sarcoma and for relapsed/refractory follicular lymphoma with EZH2 mutation^22^, and has demonstrated modest activity with durable responses in a subset of patients^23^. The variable response to EZH2 inhibition observed in preclinical studies may reflect adaptive tumor biology, including the rewiring of DNA-damage responses, metabolism, and immune signaling^24–30^, supporting rational combination strategies with EZH2 inhibitors as the path forward.

In breast cancer, EZH2 is overexpressed in the basal-like subtype, enriched for Triple-Negative Breast Cancer (TNBC), a highly aggressive subtype negative for estrogen and progesterone receptors, and HER2 amplification, which shows high relapse rates following chemotherapy^31^. Elevated *EZH2* expression in basal-like breast tumors is associated with advanced stages, chemoresistance, metastasis, and poor prognosis^32–34^. Yet, as observed in other solid malignancies, targeting EZH2 in preclinical TNBC models often yields modest effects on tumor growth despite complete loss of H3K27me^5,6,32^. This limited efficacy suggests the activation of compensatory mechanisms sustaining tumor growth, leading to tolerance to EZH2 inhibition. Understanding the molecular consequences of EZH2 inhibition in TNBC is needed to derive rational combination-based EZH2-targeting strategies^35^.

Here, we investigate the molecular consequences of targeting EZH2 in TNBC to uncover vulnerabilities that could be exploited to impair tumor growth. We show that pharmacological inhibition or depletion of EZH2 induces an integrated stress response (ISR) in TNBC cells and patient-derived xenografts (PDXs). Specifically, we demonstrate that epigenetic reprogramming induced by EZH2 inhibition activates an essential Integrated Stress Response (ISR) driven by the accumulation of misfolded proteins and double-stranded RNA (dsRNA), which activate Protein Kinase R (PKR) and Protein Kinase RNA-Like Endoplasmic Reticulum Kinase (PERK), and subsequent induction of Activating Transcription Factor-4 (ATF4). By triggering ATF4-dependent transcriptional signatures, EZH2 inhibition enhances amino acid flux and promotes glutaminolysis, in turn activating mTOR signaling and promoting glutathione-mediated antioxidant defense to support TNBC cell survival and tumor growth. Importantly, pharmacological inhibition of this metabolic adaptation with a glutaminase (GLS) inhibitor significantly enhances the efficacy of EZH2 inhibition, providing a strategy to improve TNBC response to EZH2 targeting.

## RESULTS

### EZH2 Inhibition Activates the Integrated Stress Response via PERK and PKR in TNBC

Given that EZH2 is overexpressed and associated with aggressive phenotypes in TNBC, we investigated the molecular consequences of its inhibition to identify adaptive responses and potential vulnerabilities. Consistent with prior observations^5^, treatment of TNBC PDXs (GCRC1915 and GCRC1939) with the EZH1/2 inhibitor UNC1999 (300 mg/kg, 21 days) does not significantly affect tumor growth (Fig. **1A**), despite robust reduction of H3K27me3 levels by immunoblotting and IHC (Fig. **1B-C**, **S1A-C**), and shows no induction of Cleaved Caspase-3 (Fig. **S1D-E**). In line with prior studies^36^, global H3K27ac levels, associated with accessible chromatin, are increased following EZH2 inhibition and H3K27me3 loss in TNBC PDX models. Comparable results were observed in TNBC cell lines MDA-MB-436 and Hs 578T following treatment with UNC1999 or the EZH2-selective inhibitor EPZ-6438 (1 µM, 96 h) (Fig. **1D-E** and **S1F-H**), as well as upon CRISPR/Cas9-mediated EZH2 deletion in MDA-MB-436 cells (Fig. **S1I-K**).

**Figure 1.**
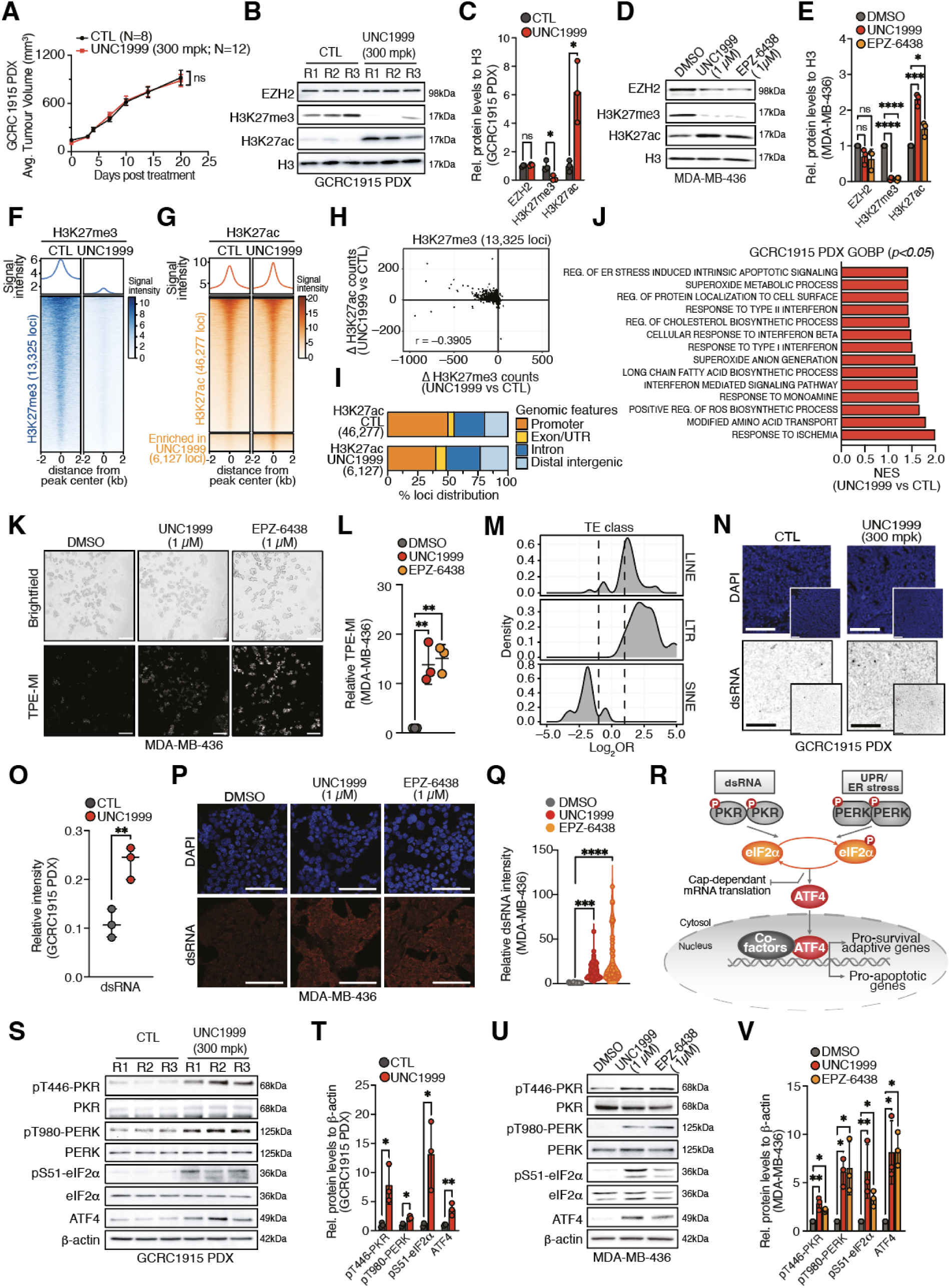
EZH2 Inhibition Induces an Integrated Stress Response Triggered by dsRNA Accumulation and Misfolded proteins in TNBC. **A.** Time-dependent growth curve showing the average tumor volume of GCRC1915 PDX implanted in female NSG mice, exposed to UNC1999 (300 mg/kg; 21 days, N=12) or control (N=8). **B.** Immunoblot showing the levels of EZH2, H3K27me3, H3K27ac, and total H3 in GCRC1915 and GCRC1939 PDX exposed or not to UNC1999 (300 mg/kg; 21 days; N=3). **C.** Quantification of the relative protein levels from Fig.1B (rel. to H3; N=3). **D.** Immunoblot showing EZH2, H3K27me3, H3K27ac, and total H3 levels in MDA-MB-436 cells exposed to UNC1999, EPZ-6438 (1 µM, 96 h; N=3) or mock treated (DMSO, 0.2%). **E.** Quantification of the relative protein levels from Fig.1D (rel. to H3; N=3). **F.** Heatmap of peak intensity signal of H3K27me3 ChIP-seq peaks (–2 to +2 kb window from peak center) upon control (left) or UNC1999 (300 mg/kg; right) treatment in GCRC1915 PDX (N=3). Compilation of tag density is illustrated for the 13,325 H3K27me3 peaks at the top of each condition. **G.** Heatmap of peak intensity signal of H3K27ac ChIP-seq specific peaks (–2 to +2 kb window from peak center) upon control (left) or UNC1999 (300 mg/kg; right) treatment in GCRC1915 PDX. Boxes illustrate common peaks between control and UNC1999, those specific to control (N=2) or UNC1999 (N=3; >2-fold difference in peak intensity between conditions, p<0.05 (DiffBind)). H3K27ac tag density plots are illustrated for the common H3K27ac peaks at the top of heapmaps for each condition. **H.** Scatter plot (Pearson, p-value<0.0001) representing the difference in H3K27ac or H3K27me3 counts on H3K27me3 sites (13,325) following UNC1999 treatment in GCRC1915 PDX (300 mg/kg, 21 days; N=2-3). **I.** Stacked bar charts illustrating the proportion of H3K27ac peaks associated with distinct genomic localization/features relative to gene TSS upon control or UNC1999 treatment in GCRC1915 PDX (300 mg/kg, 21 days; N=2-3). **J.** Gene Set Enrichment Analysis (GSEA) graph representing Normalized Enrichment Score (NES) from GOBP (Gene Ontology: Biological Process) gene set, illustrating significantly enriched pathways (p<0.05) in UNC1999-treated GCRC1915 PDX relative to control treatment (300 mg/kg; 21 days; N=3). **K.** Detection of misfolded protein load in MDA-MB-436 cells exposed to DMSO (0.2%) or EZH2 inhibitors UNC1999 or EPZ-6438 (1µM; 48 h; N=3), using the tetraphenyl ethene maleimide (TPE-MI) probe (IF). Cell density is shown in brightfield. Scale bars, 100µm. **L.** Quantification of misfolded protein load intensity from Fig.1Q showing TPE-MI intensity normalized to naïve media and relative to DMSO (0.2%). **M.** Density chart representing overall enrichment level of significantly upregulated transposable elements (TE) classes in UNC1999-treated GCRC1915 PDX (300 mg/kg; 21 days; N=3) relative to control. Odds ratio (OR) and p-value computed using Fisher’s exact test. **N.** Immunofluorescence staining using J2 (dsRNA) antibody in GCRC1915 PDX treated with control or UNC1999 (300 mg/kg; 21 days; N=3). Scale bar, 100µm. **O**. Quantification of dsRNA levels relative to DAPI intensity from Fig.1N (N=3). **P.** Immunofluorescence staining using J2-antibody in MDA-MB436 cells exposed to DMSO (0.2%), UNC1999 or EPZ-6438 (1µM; 48 h; N=3). Scale bar, 100µm. **Q.** J2-staining quantification from Fig.1P using Image J. **R.** Schematic representing the main stresses involved in the activation of the stress kinases PKR and PERK that induce ISR and ISR-mediated activation of eIF2α and ATF4 induction. **S.** Immunoblot showing pT446-PKR, PKR, pT980-PERK, PERK, pS51-eIF2α, eIF2α, ATF4 and β-actin levels in GCRC1915 PDX exposed to control or UNC1999 (300 mg/kg; 21 days; N=3). **T.** Quantification of the relative protein levels from Fig.1S (rel. to β-actin; N=3). **U.** Same as in Fig.1S in MDA-MB-436 cells exposed to DMSO (0.2%), UNC1999, or EPZ-6438 (1 µM; 48 h, N=3). **V.** Quantification of the relative protein levels from Fig.1U (rel. to β-actin; N=3). All data shown are mean ± SD for three biological replicates (C, E, L, O, Q, T and V), except for tumor volume (A) where means ± SEM and N=8 (CTL) or 12 (UNC1999) are represented. Statistical analysis was performed using one-way ANOVA (E, L, Q and V) and two-tailed Student’s t test (C, O and T). *p < 0.05, **p < 0.01, ***p < 0.001, ****p < 0.0001; ns, not significant. R: replicate #.

To investigate the epigenetic reprogramming induced by EZH2 inhibition in TNBC, we performed chromatin immunoprecipitation sequencing (ChIP-seq with exogenous *Drosophila* DNA spike-in normalization) for H3K27me3 and H3K27ac in TNBC PDX and cells. A significant decrease in the intensity and number of H3K27me3 loci is observed in GCRC1915 PDX exposed to UNC1999 in vivo (300 mg/kg, 21 days) compared to vehicle controls (Fig. **1F**; *p* < 0.0001), accompanied by a significant global increase in H3K27ac signal intensity and peak number (*p* < 0.0001) (Fig. **1G**, **S1L**). Genomic regions that lose H3K27me3 upon UNC1999 in TNBC PDXs are globally associated with increased H3K27ac (Fig. **1H**; r = -0.3905; p < 0.0001), indicating coordinated epigenetic remodeling upon EZH2 inhibition. H3K27ac peaks gained following EZH2 inhibition are preferentially enriched at distal intergenic and intronic regions (51.4% distal vs. 33% proximal-promoter), compared with untreated PDX, where H3K27ac is more promoter-proximal (44.6% distal vs. 50% proximal-promoter) (Fig. **1I**), while H3K27me3 peaks show the expected Polycomb-associated distribution across distal intergenic and intronic regions (62.1%) (Fig. **S1M**). Similar results were obtained in MDA-MB346 cells (Fig. **SN-P**). Collectively, these results reveal increased chromatin accessibility at distal non-coding regions following EZH2 inhibition in TNBC.

The persistence of TNBC growth despite extensive epigenetic rewiring prompted us to assessed the functional consequences of this epigenetic shift. We analyzed transcriptional changes following EZH2 inhibition in TNBC PDX models and cell lines. Gene set enrichment analysis (GSEA) of genes upregulated following UNC1999 treatment in TNBC PDXs shows significant enrichment of stress-associated pathways (Gene Ontology (GO) Biological Pathways), including ischemia, amino acid transport, ROS and superoxide metabolism, interferon signaling, cholesterol and fatty acid biosynthesis, and endoplasmic reticulum (ER) stress (Fig. **1J**). We next assessed whether some of these stresses are induced upon EZH2 inhibition in TNBC. Using the tetraphenylethene maleimide (TPE-MI) thiol probe, which detects free cysteine thiols, we observe the accumulation of misfolded proteins, a hallmark of endoplasmic reticulum (ER) stress, following exposure to UNC1999 or EPZ-6438 in TNBC cells (Fig. **1K-L** and **S2A-B**). We also observe that EZH2 inhibition induces the production of double-stranded RNA (dsRNA), a hallmark of interferon signaling, a stress-responsive innate immune pathway. Indeed, RNA-seq reveals increased expression of non-coding transposable elements (TEs) in UNC1999-treated TNBC PDXs, particularly Long Terminal Repeat (LTR; ERV1, ERVL/K) and LINE-1 (L1) families (Fig. **1M**, **S2C**). These TEs, which can form dsRNA upon derepression, are enriched in intergenic/intronic non-coding regions that gain accessibility following EZH2 inhibition in TNBC PDX (Fig. **S2D**). Accordingly, dsRNA accumulation detected by immunofluorescence using the J2 antibody is significantly increased following EZH2 inhibition in TNBC PDXs (Fig. **1N-O** and **S2E;** UNC1999, 300 mg/kg) and MDA-MB436 cells (Fig. **1P-Q**; UNC1999/EPZ-6438, 1 µM, 48 h). Together, these observations indicate that EZH2 inhibition in TNBC triggers epigenetic rewiring that induces cellular stresses, including misfolded protein and dsRNA accumulation.

Accumulation of dsRNA is known to activate Protein Kinase R (PKR) signaling, while misfolded proteins are canonical triggers of the PKR-like endoplasmic reticulum kinase (PERK) arm of the unfolded protein response (UPR) (Fig. **1R**). PERK and PKR phosphorylate the eukaryotic initiation factor 2α (eIF2α) to activate the integrated stress response (ISR), a conserved signaling pathway that reprograms translation and gene expression. This includes preferential translation of Activating Transcription Factor 4 (ATF4), which promotes restoration of cellular homeostasis or adaptive survival under stress. Consistent with this, EZH2 inhibition induces activation of PKR (pT446-PKR), PERK (pT980-PERK), and eIF2α (pS51-eIF2α), along with robust induction of ATF4, in TNBC PDX models (GCRC1915, GCRC1939; Fig. **1S-T** and **S2F-H**; UNC1999, 300 mg/kg) and in TNBC cell lines, including MDA-MB-436 (Fig. **1U-V**; UNC1999 or EPZ-6438, 1 µM, 48 h) and Hs 578T (Fig. **S2I-J**), as assessed by immunoblotting or IHC. Induction of ATF4 is also observed in EZH2-KO MDA-MB-436 cells compared to sgCtl cells (Fig. **S2K-L**). Together, these data establish that EZH2 inhibition drives extensive epigenetic reprogramming in TNBC, triggering PERK- and PKR-dependent activation of the integrated stress response.

### The Activation Integrated Stress Response Supports TNBC Cell Survival upon EZH2 Inhibition

Since EZH2 inhibition activates stress responses without impairing TNBC growth, we next assessed the functional role of ISR induction by depleting core ISR components with independent siRNAs in MDA-MB-436 and Hs578T cells (Fig. **2A**, **S2M**). Depletion of PERK, PKR, or eIF2α had minimal effects on basal cell viability in TNBC cells but significantly reduced cell viability specifically upon UNC1999 treatment (1 µM, 96 h) (Fig. **2B** and **S2N**). Similarly, siRNA-mediated depletion of ATF4 modestly reduced viability under basal conditions (10–25%) but resulted in a profound loss of viability (80–95%) and robust induction of the apoptotic marker cleaved PARP (cl-PARP) upon EZH2 inhibition in MDA-MB-436, Hs578T (Fig. **2C-E**; UNC1999, 1 µM, 96 h). Similar results were observed with GCRC1915 patient-derived cells exposed to UNC1999 (1 µM; Fig. **2F-G**). Consistent with these genetic perturbations, pharmacologic restoration of eIF2B activity using the Integrated Stress Response Inhibitor (ISRIB; 100 nM), which counteracts the inhibitory effects of pS51-eIF2α, significantly sensitized TNBC cells to EZH2 inhibition (Fig. **2H**), inducing the apoptotic marker cl-PARP (Fig. **2I-J**; UNC1999, 1 µM, 96 h). Collectively, these findings demonstrate that ISR signaling induction supports TNBC cell survival under EZH2 inhibition.

**Figure 2.**
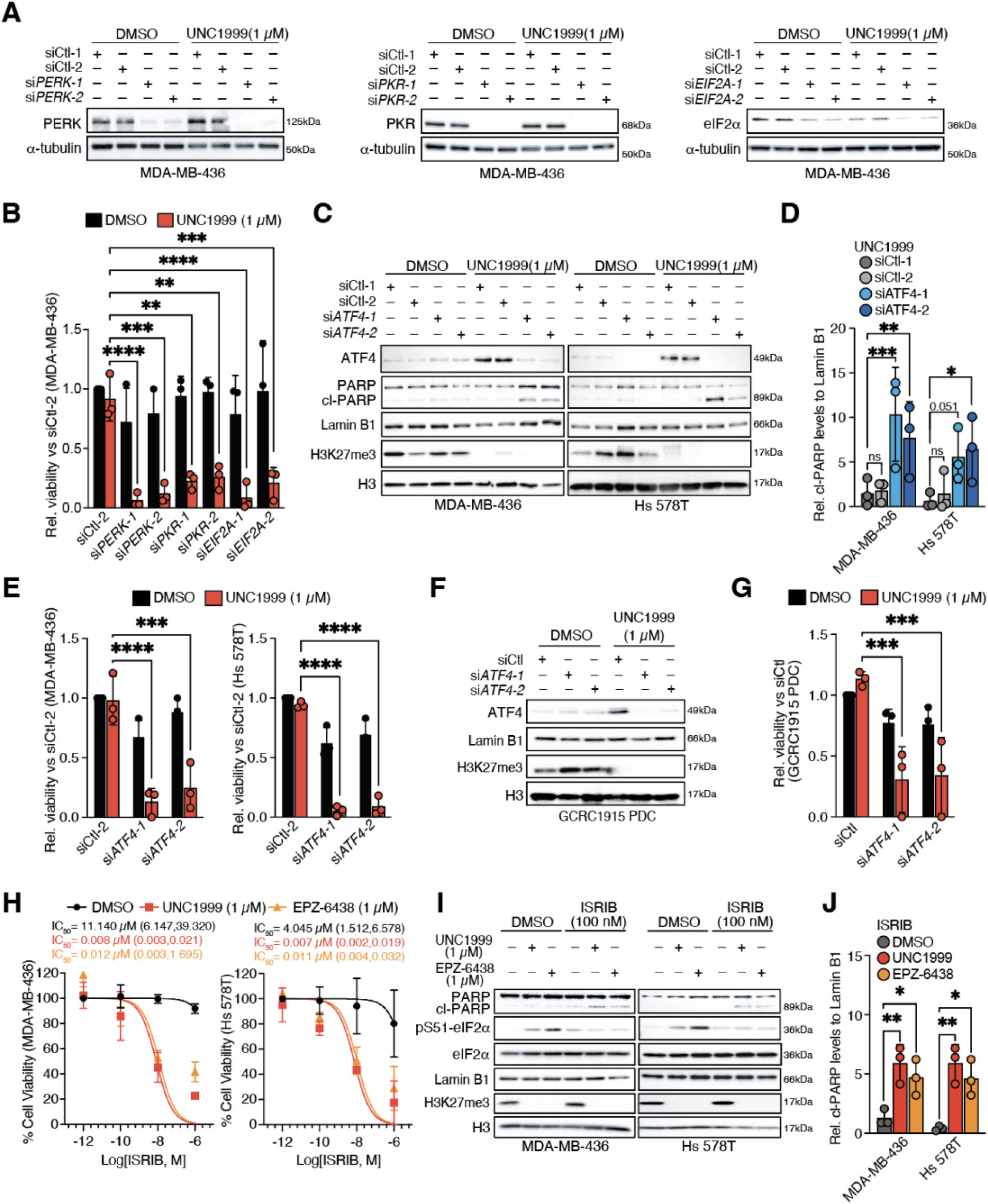
Induction of the Integrated Stress Response Supports TNBC Cell Survival upon EZH2 Inhibition. **A.** Immunoblot showing the levels of the indicated proteins (PKR, PERK, eIF2α or α-tubulin) in MDA-MB-436 cells exposed to DMSO (0.2%) or UNC1999 (1µM; 48 h, N=3) and depleted or not for the indicated protein using two siRNAs per gene (vs non-targeting siRNAs (siCtl)). **B.** Relative viability (crystal violet) of MDA-MB-436 cells depleted of either PKR, PERK, or eIF2α using siRNAs (vs siControl) and exposed to UNC1999 (1µM; 96 h, N=3) or DMSO (0.2%). **C.** Immunoblot showing the levels of the indicated proteins in MDA-MB-436 and Hs 578T cells exposed to DMSO (0.2%) or UNC1999 (1µM; 72 h; N=3) and depleted of ATF4 using two siRNAs (vs siCtl). **D.** Quantification of the relative protein levels from Fig. 2C (rel. to Lamin B1; N=3). **E.** Relative viability (crystal violet) of MDA-MB-436 and Hs 578T cells depleted of ATF4 using siRNAs (vs siCtl) and exposed to UNC1999 (1µM; 96 h) or DMSO (0.2%; N=3). **F.** Immunoblot showing the levels of the ATF4, Lamin B1, H3K27me3 and total H3 proteins in GCRC1915 PDC exposed to UNC1999 (1µM; 48 h) or DMSO (0.2%; N=3) and depleted of ATF4 using specific siRNAs (72 h). **G.** Relative viability (crystal violet) of GCRC1915-PDC depleted of ATF4 using 2 siRNAs and exposed to UNC1999 (1µM; 96 h; N=3). **H.** Dose-response curves representing the relative normalized viability of MDA-MB-436 (left) and Hs 578T (right) cells exposed to DMSO (0.2%), UNC1999 or EPZ-6438 (1µM; 96h) (N=3) and co-treated with varying doses of ISRIB. IC50 (95% Confidence Interval) was calculated and indicated at the top of each graph. **I.** Immunoblot showing the levels of the indicated proteins in MDA-MB-436 and Hs 578T cells exposed to DMSO (0.2%), UNC1999 and EPZ-6438 (1µM; 72 h; N=3) and treated with the integrated stress response inhibitor (ISRIB; 100nM; vs DMSO). **J.** Quantification of the relative protein levels from Fig. 2H (rel. to Lamin B1; N=3). Data are the mean ± SD from 3 biological replicates (B, D, E, G, H and J). Statistical analysis performed using one-way (D and J) and two-way ANOVA (B, E and G). ns, not significant, *p < 0.05, **p < 0.01, ***p < 0.001, ****p < 0.0001. R: replicate #.

### EZH2 inhibition induces an ATF4-dependent Amino Acid Metabolic Transcriptional Program in TNBC

Given that ISR activation supports TNBC survival upon EZH2 inhibition, we next investigated the downstream mechanisms by which this stress program promotes cell viability. GSEA of differentially expressed genes in GCRC1915 PDX and MDA-MB-436 cells exposed to UNC1999 had revealed significant enrichment of previously defined ATF4 target genes, including amino acid transporters and metabolic enzymes implicated in redox homeostasis (Fig. **1J** and **S3A**). Consistent with this enrichment, EZH2 inhibition using UNC1999 in GCRC1915 PDX models (300 mg/kg) significantly increased the expression of genes enriched in amino acid transport, antioxidant defense, and amino acid metabolism (Fig. **3A** and **S3B**). Immunoblotting and immunohistochemical analyses confirmed the significant induction of key amino acid transporters (LAT1/SLC7A5, CD98/SLC3A2, ASCT2/SLC1A5, xCT/SLC7A11) and metabolic enzymes (PSAT1, BCAT1, ASNS) at the protein level following EZH2 inhibition in both GCRC1915 PDX and TNBC cell lines (Fig. **3B-E** and **S3C-E**). To determine whether this response depends on induction of ATF4 in response to EZH2 inhibition, we depleted ATF4 using independent siRNAs in MDA-MB-436 cells. We observed that loss of ATF4 significantly attenuated the UNC1999-dependent induction of these transporters and metabolic enzymes in TNBC cells (Fig. **3F-G** and **S3F-G**).

**Figure 3.**
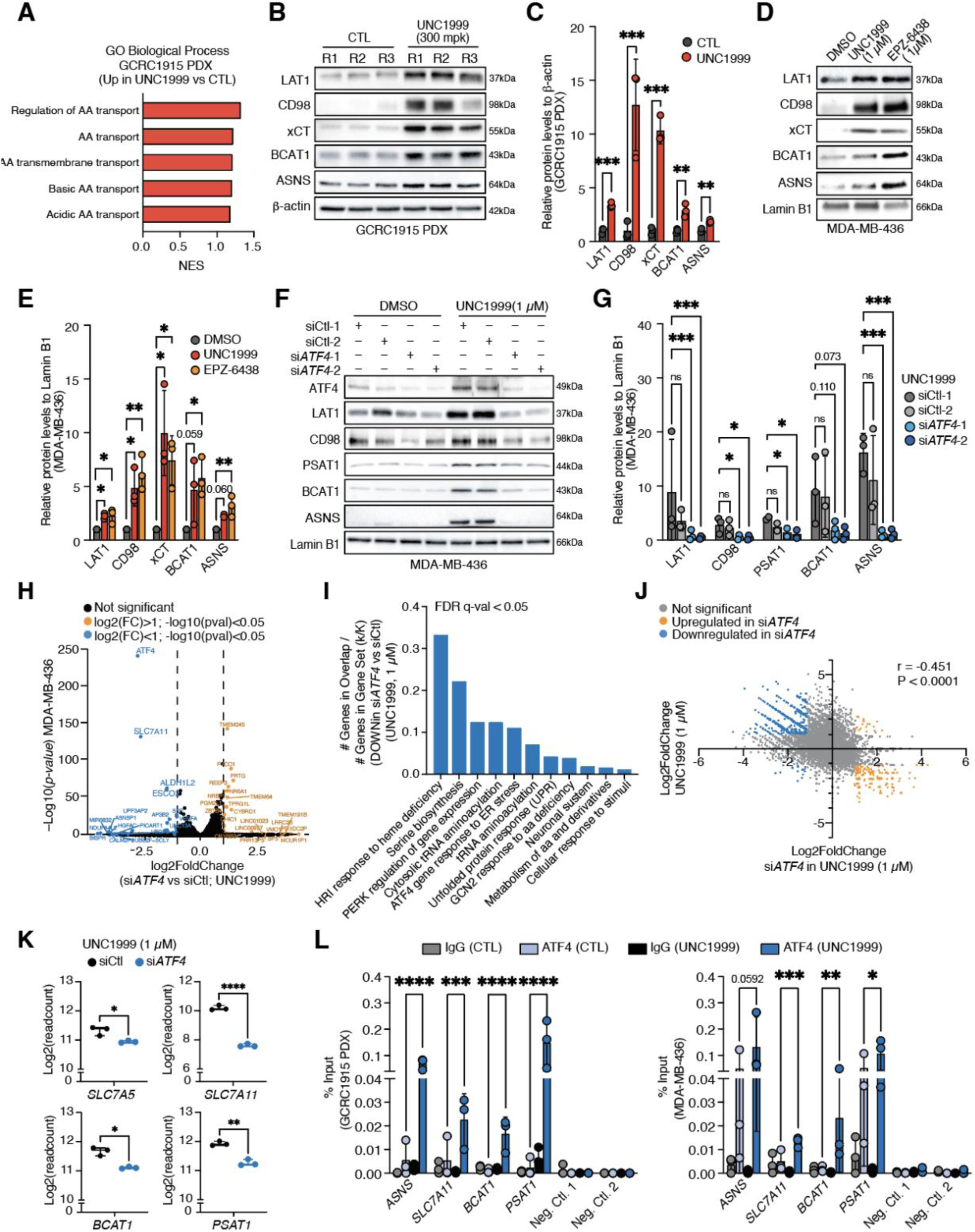
ATF4 Regulates an Amino Acid Gene Signature in Response to EZH2 Inhibition in TNBC. **A.** GSEA bar graph displaying significantly enriched metabolism-related gene ontology biological process (GOBP) pathways (p<0.05), from genes significantly up-regulated in UNC1999-treated GCRC1915-PDX (300 mg/kg; 21 days) relative to control (N=3). **B.** Immunoblot showing LAT1, CD98, xCT, BCAT1, ASNS and β-actin levels in GCRC1915 PDX exposed to control or UNC1999 (300 mg/kg; 21 days; N=3). **C.** Quantification of the relative protein levels from Fig. 3B (rel. to β-actin; N=3). **D.** Same as in B in MDA-MB436 cells exposed to DMSO (0.2%) or UNC1999 (1µM; 96 h; N=3). **E.** Quantification of the relative protein levels from Fig. 3D (rel. to Lamin B1; N=3). **F.** Immunoblot showing the levels of ATF4, LAT1, CD98, PSAT1, BCAT1, ASNS and Lamin B1 in MDA-MB-436 cells exposed to DMSO (0.2%) or UNC1999 (1µM; 72 h; N=3) and depleted for ATF4 using two siRNAs against *ATF4* (72 h) vs non-targeting siRNA (siCtl). **G.** Quantification of the relative protein levels from Fig. 3F (rel. to Lamin B1; N=3). **H.** Volcano plots showing the -Log10(*p-value*) and the Log2 fold change in gene expression upon ATF4 depletion by siRNA (48 h) in MDA-MB-436 cells exposed to UNC1999 (1μM, 48 h, N=3). **I.** Bar graph representing up-regulated pathways (GSEA) following ATF4 depletion in MDA-MB-436 cells exposed to UNC1999 (1µM; 48 h; N=3) vs DMSO (0.2%). **J.** Scatter plot representation of Log2 fold change in gene expression resulting from UNC1999 vs DMSO (vertical) and si*ATF4* vs siCtl in UNC1999-treated MDA-MB-436 cells (1µM; 48 h; N=3) (r= -0.451; p-value<0.0001). **K.** Gene expression levels of *SLC5A7*, *SLC7A11*, *PSAT1,* and *BCAT1* upon depletion of ATF4 (siATF4) vs siCtl in MDA-MB-436 cells exposed to UNC1999 (1µM; 48 h; N=3). **L.** ATF4 Chromatin immunoprecipitation followed by qPCR amplification at specific ATF4 target gene promoters (*ASNS*, *SLC7A11*, *BCAT1,* and *PSAT1*) in none-treated vs UNC1999-treated GCRC1915 PDX (left; 300 mg/kg; 21 days; N=3) and MDA-MB-436 (right; 1µM; 48 h). Data are shown as %input. The data presented are the mean ± SD from 3 biological replicates (C, E, G, K and L). Statistical analysis was performed using one-way ANOVA (E, G and L) and two-tailed Student’s t test (C and K). ns, not significant, *p < 0.05, **p < 0.01, ***p < 0.001, ****p < 0.0001. R: replicate #.

Transcriptomic profiling further demonstrated that genes downregulated upon ATF4 depletion in EZH2-inhibited cells are significantly enriched for ISR-related pathways, including PERK and amino acid metabolism (Reactome, FDR q < 0.05; Fig. **3H-I**). Notably, genes induced by UNC1999 treatment in TNBC cells show a significant inverse correlation with gene expression changes following ATF4 depletion (Fig. **3J**), including multiple amino acid transporters and metabolic enzymes involved in redox homeostasis (Fig. **3K**). In contrast, ATF4 depletion leads to upregulation of gene signatures associated with senescence, apoptosis, oxidative stress, and ferroptosis (Fig. **S3H**), consistent with a protective role for ATF4 under EZH2 inhibitory stress. Finally, chromatin immunoprecipitation followed by qPCR demonstrates that EZH2 inhibition promotes ATF4 recruitment to the promoters of representative amino acid transport and metabolism genes, including *ASNS*, *BCAT1*, *PSAT1*, and *SLC7A11*, in GCRC1915 PDX and MDA-MB-436 cells (Fig. **3L**), supporting a direct transcriptional regulatory role. Together, these results demonstrate that EZH2 inhibition engages ATF4 to induce a transcriptional program centered on amino acid metabolism and redox homeostasis in TNBC.

### EZH2 Inhibition Activates an Amino Acid-Dependent Metabolic Rewiring in TNBC

Given the ATF4-dependent induction of amino acid transport and metabolic genes following EZH2 inhibition, we next assessed whether this program translates into functional changes in amino acid handling in TNBC cells. Quantification of metabolites in conditioned media from MDA-MB-436 and Hs 578T cells revealed a significant increase in glutamine uptake and glutamate release following treatment with UNC1999 or EPZ-6438 (1µM; 72 h), while glucose uptake and lactate secretion remained unchanged (Fig. **4A** and Fig. **S4A**; Nova BioProfile 400 analyzer). Targeted amino acid profiling using LC-QQQ-MS further demonstrated that EZH2 inhibition enhances uptake of branched-chain amino acids (leucine, valine, isoleucine), serine, methionine, glutamine, and cystine, accompanied by increased glutamate release (Fig. **4B** and Fig. **S4B**). These metabolic changes are consistent with the engagement of amino acid transporters, including LAT1 and the xCT/CD98 antiporter system, which are upregulated following EZH2 inhibition (Fig. 3A-B). The xCT/CD98 complex couples cystine import to glutamate export, thereby supporting glutathione synthesis and antioxidant defense. In line with this model, uptake of glutamine and branched-chain amino acids can replenish intracellular glutamate through transamination reactions, while serine and methionine contribute to glutathione biosynthesis and redox-related metabolic pathways (Fig. **4C**). Importantly, depletion of ATF4 significantly attenuated EZH2 inhibitor-induced amino acid uptake in MDA-MB-436 cells, indicating that these metabolic changes are ATF4-dependent (Fig. **4D**).

**Figure 4.**
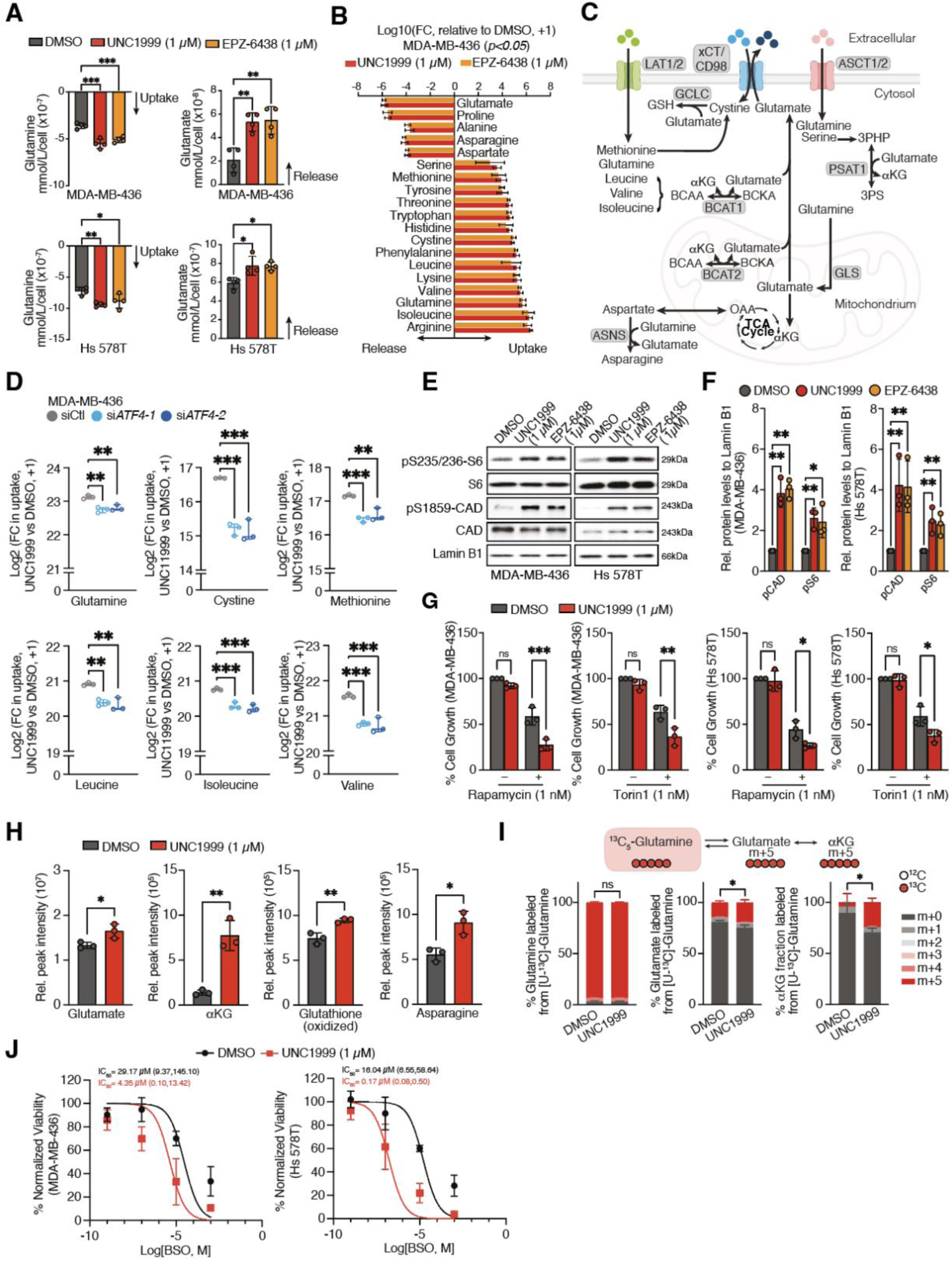
EZH2 Inhibition Induces ATF4-Mediated Metabolic Adaptation in TNBC. **A.** Relative quantification of glutamine and glutamate (BioProfile 400, Nova Biomedical) in conditioned media exposed to MDA-MB-436 or Hs 578T cells treated with UNC1999 or EPZ-6438 (1µM; 72 h) or DMSO (0.2%; N=4). **B.** Relative metabolite quantification in conditioned media using Triple Quadrupole-Liquid Chromatography-Mass Spectrometry (QQQ-LC-MS; Intrada-amino-acids) in MDA-MB-436 cells exposed to UNC1999 or EPZ-6438 (p<0.05; 1µM; 72 h) or DMSO (0.2%; N=4). Metabolite levels for each experimental condition are normalized to those of the naïve medium. **C.** Schematic representation of amino acid uptake and glutaminolysis coupled with transaminase reactions and antioxidant response. **D.** Relative metabolite levels in conditioned media using QQQ-LC-MS (Intrada-amino-acids) in MDA-MB-436 cells exposed to DMSO (0.2%) or UNC1999 (1µM; 60h) and transfected with non-targeting siRNA (siCtl) or two siRNA against *ATF4* (20nM; 60h; N=3). Metabolite levels for each experimental condition are normalized against those of the naïve medium. **E.** Immunoblot showing pS1859-CAD, CAD, pS235/236-S6, S6 and Lamin B1 levels in MDA-MB-436 or Hs 578T cells exposed to UNC1999, EPZ-6438 (1µM; 96 h) or DMSO (0.2%; N=3). **F.** Quantification of the relative protein levels from Fig. 4D (rel. to Lamin B1; N=3). **G.** Percent growth of MDA-MB-436 and Hs 578T cells exposed to UNC1999 (1µM; 96 h) or DMSO (0.2%; N=3) exposed to DMSO (0.1%), Rapamycin (1nM) or Torin1 (1nM). **H.** Quantification of selected intracellular metabolites in Hs 578T cells using Quadrupole Time-of-Flight (QTOF)-LC-MS (Ion-pairing) exposed to UNC1999 (1µM; 72 h) or DMSO (0.2%; N=3). **I.** Stack bars illustrating stable isotope tracing analysis (SITA) using glutamine-deprived media supplemented with ^13^C_5_-glutamine (4mM) to identify the fractional pool of labeled glutaminolysis intermediates (glutamine m+5, glutamate m+5 and ⍺KG m+5) in Hs 578T cells exposed to DMSO (0.2%) or UNC1999 (1μM, 96 h), quantified by QTOF-LC-MS (ion pairing) after 30 minutes exposure to ^13^C_5_-glutamine. **J.** Dose-response curves representing the relative normalized viability of MDA-MB-436 (left) and Hs 578T (right) cells exposed to DMSO (0.2%) or UNC1999 (1µM; 96h; N=3) and co-treated with varying doses of L-Buthionine sulfoximine (BSO), a potent inhibitor of the catalytic subunit of glutamate-cysteine ligase (GCLC). IC50 (95% Confidence Interval) was calculated and indicated at the top of each graph. The data shown are the mean ± SD from 3 to 4 biological replicates (A, B, D, F, G, H, I and J). Statistical analysis was performed by one-way (A, D and F) and two-way ANOVA (G), and two-tailed Student’s t test (H and I). ns, not significant, *p < 0.05, **p < 0.01, ***p < 0.001.

Because the mammalian/mechanistic target of rapamycin (mTOR) complex 1 (mTORC1) signaling is highly sensitive to intracellular amino acid availability, we next evaluated its activation following EZH2 inhibition in TNBC cells and observed increased phosphorylation of the downstream Ribosomal Protein S6 Kinase (S6K) target S6 (pS235/236-S6; Fig. **4E-F**). Consistent with mTORC1 activation, EZH2 inhibition increased phosphorylation of the direct S6K1 substrate carbamoyl-phosphate synthetase 2, aspartate transcarbamylase, and dihydroorotase enzyme (CAD, pS1859-CAD), which couples mTORC1-driven amino acid sensing to de novo pyrimidine synthesis. Furthermore, EZH2 inhibition by UNC1999 (1µM) further impaired MDA-MB436 and Hs 578T cell growth in combination with pharmacological inhibition of mTOR signaling using rapamycin (mTORC1-selective) or Torin1 (dual mTORC1/2 inhibitor), indicating that mTOR signaling contributes to growth-supportive response upon EZH2 inhibition in TNBC cells (Fig. **4G**).

LC-QTOF-MS analysis further reveals increased intracellular levels of glutamate, α-ketoglutarate (αKG), oxidized glutathione, asparagine, fumarate, and malate in TNBC cells treated with UNC1999 (1 µM, 72 h) (Fig. **4H** and Fig. **S4C**), consistent with changes in amino acid uptake and utilization and engagement of redox-associated metabolic pathways. Stable isotope tracing with ^13^C_5_-glutamine demonstrates significantly increased incorporation of glutamine-derived carbon into glutamate and α-ketoglutarate upon EZH2 inhibition in TNBC cells, indicative of enhanced glutaminolysis, while carbon flux into the oxidative branch of the TCA cycle is modestly reduced with a corresponding increase in reductive carboxylation-derived metabolites (Fig. **4I** and Fig. **S4D-E**). No change in glutamine-derived labeling of GABA is observed, indicating that this alternative anaplerotic route is not engaged (Fig. **S4F**). Despite these metabolic changes, Seahorse analysis revealed no significant alterations in basal or stress-induced oxygen consumption or extracellular acidification rates (OCR, ECAR), suggesting that EZH2 inhibition does not induce broad bioenergetic rewiring in TNBC cells (Fig. **S4G-H**).

Given the increased cystine uptake, enhanced glutaminolysis, glutamate availability and release, and accumulation of oxidized glutathione following EZH2 inhibition, we next tested whether TNBC cells become functionally dependent on glutathione-dependent redox homeostasis under these conditions. Inhibition of the glutathione-synthesizing enzyme glutamate–cysteine ligase with buthionine sulfoximine (BSO) significantly reduced TNBC cell viability, specifically when treated with UNC1999 (1 µM), revealing a metabolic vulnerability associated with induced amino acid-and redox-related adaptations upon EZH2 inhibition (Fig. **4J**). Together, these results demonstrate that EZH2 inhibition engages an ATF4-dependent amino acid metabolic program that activates mTOR signaling and redox-maintenance, thereby supporting TNBC cell growth and fitness under EZH2-inhibitory stress.

### Glutaminase Targeting Sensitizes TNBC to EZH2 Inhibition

To determine whether the observed enhancement of glutamine metabolism is functionally required under EZH2 inhibition, we assessed TNBC cell viability under glutamine-replete or glutamine-limited conditions. EZH2 inhibition with UNC1999 (1 µM) selectively reduces cell viability under low-glutamine conditions (0.08 mM), while cells remain relatively insensitive in glutamine-replete media (4 mM), indicating increased glutamine dependency upon EZH2 inhibition (Fig. **5A**). Consistent with this phenotype, EZH2 inhibition using UNC1999 or EPZ-6438 (1µM) significantly increases the expression of glutaminase (GLS), the enzyme catalyzing the first step of glutaminolysis, in TNBC cell lines (Fig. **5B-C**). GLS levels are also elevated in EZH2-deleted TNBC cells (Fig. **S5A-B**) and in TNBC PDX models (GCRC1915 and GCRC1939) treated with UNC1999, as assessed by immunoblotting and immunohistochemistry (Fig. **5D-E** and Fig. **S5C**). Importantly, independent siRNA-mediated depletion of ATF4 significantly blunts UNC1999-induced GLS upregulation in TNBC cells, indicating that ATF4 induction contributes to GLS increase downstream of EZH2 inhibition (Fig. **5F-G**).

**Figure 5.**
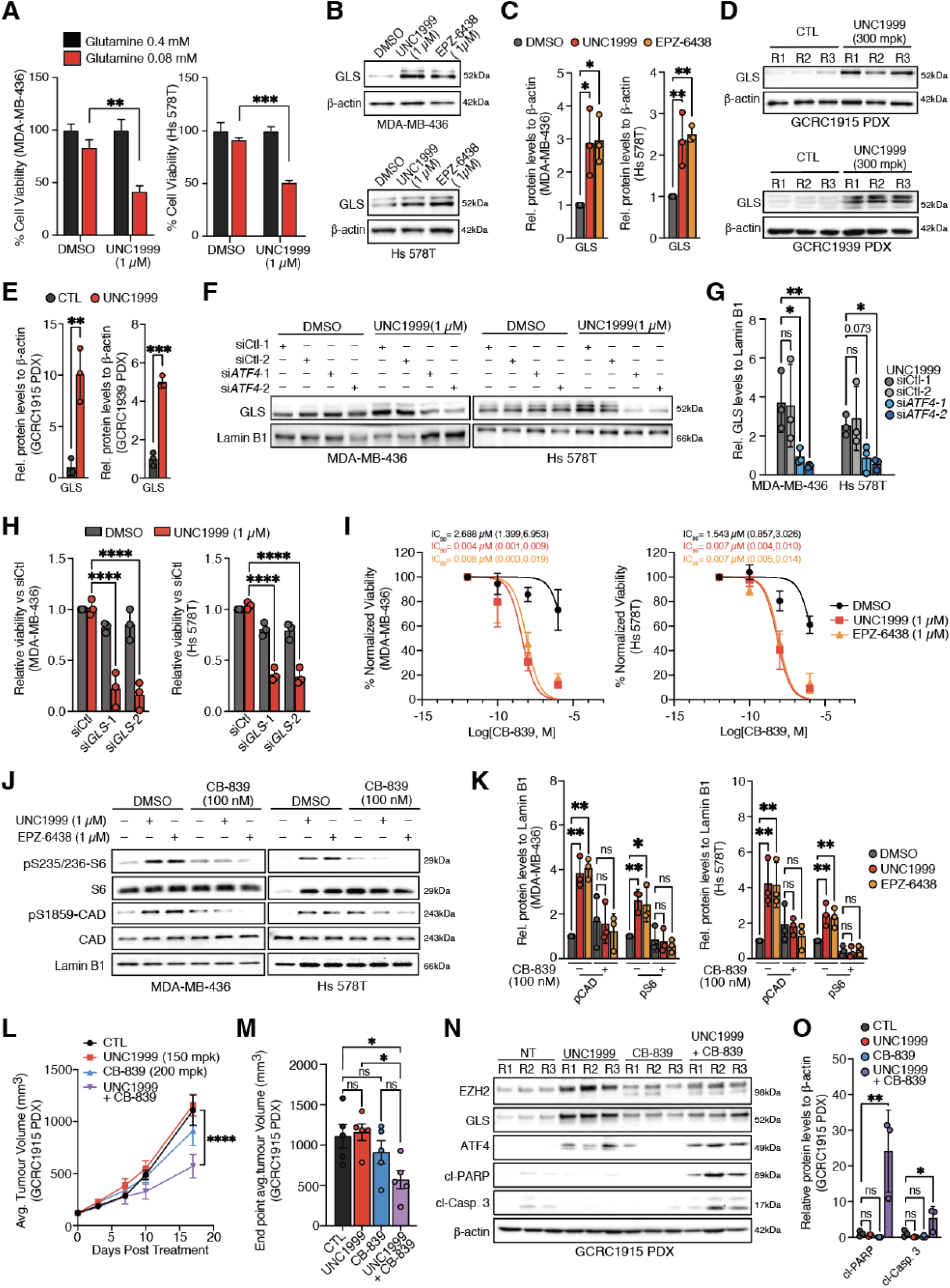
Blocking glutaminase sensitizes TNBC to EZH2 inhibition. **A.** Percent viability of MDA-MB-436 (left) or Hs 578T (right) cells exposed to UNC1999 (1µM; 96 h) or DMSO (0.2%; N=3) exposed to either full media (0.4 mM glutamine) or to media depleted in glutamine (0.08 mM) (data shown relative to full media). **B.** Immunoblot showing GLS and β-actin levels in MDA-MB-436 or Hs 578T cells exposed to UNC1999, EPZ-6438 (1µM; 96 h) or DMSO (0.2%; N=3). **C.** Quantification of the relative GLS levels from Fig. 5B (rel. to β-actin; N=3). **D.** Immunoblot showing GLS and β-actin levels in GCRC1915 and GCRC1939 PDXs exposed or not to UNC1999 (300 mg/kg; 21 days; N=3). **E.** Quantification of the relative GLS levels from Fig. 5D (rel. to β-actin; N=3). **F.** Immunoblot showing GLS and Lamin B1 levels in MDA-MB-436 or Hs 578T cells exposed to UNC1999 (1µM; 72 h) or DMSO (0.2%; N=3) and depleted of ATF4 using two siRNAs against *ATF4* (60h) vs non-targeting siRNAs (siCtl). **G.** Quantification of the relative GLS levels from Fig. 5F (rel. to Lamin B1; N=3). **H.** Relative viability (crystal violet) of MDA-MB-436 and Hs 578T cells depleted of GLS using siRNAs (vs siCtl) and exposed to UNC1999 (1µM; 96 h) or DMSO (0.2%; N=3). **I.** Dose-response curves representing the relative normalized viability of MDA-MB-436 (left) and Hs 578T (right) cells exposed to DMSO (0.2%), UNC1999 or EPZ-6438 (1µM; 120h) (N=3) and co-treated with varying doses of CB-839. IC50 (95% Confidence Interval) was calculated and indicated at the top of each graph. **J.** Immunoblot showing pS1859-CAD, CAD, pS235/236-S6, S6 and Lamin B1 levels in MDA-MB-436 or Hs 578T cells treated with DMSO (0.2%), UNC1999 or EPZ-6438 (1µM; 96 h; N=3) and exposed to CB-839 (100nM). **K.** Quantification of the relative protein levels from Fig. 5J (rel. to Lamin B1; N=3). **L.** Graph showing average tumor volume over time for GCRC1915 PDX implanted in female NSG mice and treated as indicated (control, CB-839 (200 mg/kg; 17 days), UNC1999 (150 mg/kg; 17 days) or combination) (17 days; N=5 per group). **M.** Bar graph showing average tumor volume as in J on day 17 (endpoint). **N.** Immunoblot showing EZH2, GLS, ATF4, cl-PARP, cl-Caspase3, and β-actin levels in GCRC1915 PDX tumors treated with CTL, CB-839 (200 mg/kg), UNC1999 (150 mg/kg), or combination (17 days; N=3). **O.** Quantification of the relative cl-PARP and cl-caspase3 levels from Fig. 5N (rel. to β-actin; N=3). Data represented are means ± SD from 3 biological replicates (A, C, E, G, H, I, K and O); error bars represent SEM for tumor volume (L and M). Statistical analysis was performed by one-way (C, G, K, M and O) and two-way ANOVA (H and L) and two-tailed Student’s t test (A and E). ns, not significant, *p < 0.05, **p < 0.01, ***p < 0.001, ****p < 0.0001. R: replicate #.

To directly assess the functional contribution of GLS in the response to EZH2 inhibition, we depleted GLS using independent siRNAs (Fig. **5H** and **S5D**) and observed a pronounced sensitization of TNBC cells to EZH2 inhibition (UNC1999/EPZ-6438, 1 µM, 120 h). Likewise, pharmacological inhibition of GLS using the clinically relevant inhibitor telaglenastat (CB-839) similarly sensitized multiple TNBC cell lines and patient-derived cells to EZH2 inhibition (Fig. **5I** and Fig. **S5E** UNC1999/EPZ-6438, 1 µM, 120 h), as well as EZH2-deficient cells (Fig. **S5F**), indicating that loss of EZH2 function creates a dependency on glutaminase activity. Mechanistically, GLS inhibition attenuated EZH2 inhibitor-induced activation of mTOR signaling, as evidenced by reduced phosphorylation of pS235/236-S6 and pS1859-CAD, supporting a model in which enhanced glutaminolysis sustains mTOR-dependent growth signaling following EZH2 inhibition (Fig. **5J-K**).

Finally, to evaluate its therapeutic relevance in vivo, we treated mice bearing GCRC1915 TNBC PDX tumors with UNC1999 (150 mpk), CB-839 (200 mpk), or the combination. Combined EZH2 and GLS inhibition significantly reduced tumor growth compared to single-agent treatments, without affecting body weight (Fig. **5L-M** and Fig. **S5G**). Tumor analyses revealed robust induction of apoptotic markers, including cleaved PARP and cleaved caspase-3, specifically in the combination-treated group (Fig. **5N-P**), with no corresponnding increase in apoptosis or stress markers observed in normal tissues (Fig. **S5H**). Similar sensitization to GLS inhibition was observed in xenografts derived from EZH2-deleted TNBC cells (Fig. **S5I-L**). Together, these findings demonstrate that EZH2 inhibition induces an ATF4-dependent glutamine metabolic rewiring that creates a selective dependency on glutaminase activity and that targeting this vulnerability markedly sensitizes TNBC cells and tumors to EZH2 inhibition.

## DISCUSSION

This study defines a coordinated adaptive response of TNBC cells to EZH2 inhibition, linking epigenetic perturbation to cellular stress signaling and metabolic reprogramming. Although EZH2 inhibition induces widespread epigenetic changes and elicits multiple stress responses that are potentially deleterious to cancer cells, including misfolded proteins and dsRNA accumulation, it does not uniformly impair TNBC cell viability, proliferation, or tumor growth. Instead, we show that TNBC cells adapt to EZH2-inhibitory stress by engaging an ISR that supports cellular fitness and proliferative capacity. Central to this response is ISR-driven ATF4 induction, which promotes a metabolic rewiring characterized by enhanced amino acid uptake and glutaminolysis, thereby supporting glutathione-dependent redox homeostasis and mTOR activation to sustain TNBC cell growth upon EZH2 inhibition. Importantly, we demonstrate that disrupting this adaptive metabolic response, specifically by targeting glutaminolysis through GLS inhibition, selectively sensitizes TNBC cells and tumors to EZH2 inhibition, revealing a therapeutically exploitable vulnerability.

EZH2 is emerging as a promising therapeutic target in cancer due to its central role in regulating gene expression and cell identity through histone modifications. It has been implicated in cell proliferation, invasion, migration, drug resistance and senescence, underscoring its significance in tumor progression^25,37,38^. The development of pharmacological EZH2 inhibitors has therefore garnered considerable attention, as exemplified by the FDA approval of tazemetostat (TAZVERIK®) for epithelioid sarcoma and follicular lymphoma^22^. Recent studies have highlighted the context-dependent nature of EZH2 targeting, as its inhibition does not uniformly suppress tumor growth and, in certain settings, may even promote tumorigenesis^39^. The tumor microenvironment further modulates EZH2 activity, with hypoxia and inflammation regulating EZH2 expression through HIF-1α^40^ and NF-κB/STAT3^41^ signaling pathways, while nutrient availability can influence its activity via metabolic changes^42^. Adding another layer of complexity, EZH2 also exhibits PRC2-independent functions that have been described in TNBC and may persist despite pharmacological inhibition, thereby contributing to tumorigenesis through alternative mechanisms^43^. Whether such non-canonical EZH2 activities contribute to the ISR-ATF4-driven metabolic reprogramming identified in this study remains an important question for future investigation. Collectively, these observations underscore the need for a comprehensive understanding of the multifaceted and context-dependent roles of EZH2 across cancer types to optimize therapeutic strategies and minimize unintended consequences.

EZH2 inhibition induces broad epigenetic remodeling in TNBC, characterized by loss of H3K27me3 and compensatory gains in H3K27ac at promoters and intergenic regions, consistent with prior reports linking this epigenetic switch to adaptive transcriptional programs that activate oncogenic pathways^35,44^. In our models, this remodeling is accompanied by misfolded protein accumulation, ER stress signatures, as well as increased expression of transposable elements, dsRNA and induction of innate immune-associated stress signatures, including interferon-related pathways. In some tumor contexts, induction of this signature can potentiate the antitumor activity of EZH2 inhibition^45^ and enhance sensitivity to immunomodulatory therapies when amplified^46–48^. However, these responses are not sufficient to impair tumor growth in TNBC. Instead, our data indicate that these epigenetic stress outputs act as upstream stress signals that converge on ISR activation, which ultimately determines cellular fitness following EZH2 inhibition. This framework helps reconcile why EZH2 inhibitors can elicit ER stress and dsRNA or immune-related transcriptional programs without uniformly inducing cell-intrinsic cytotoxicity and suggests that epigenetic stress alone is insufficient to drive tumor regression in the absence of targeted disruption of adaptive stress-response pathways. Targeting compensatory epigenetic adaptations has emerged as an effective strategy to enhance responses to EZH2 inhibition^49^. Consistent with this concept, co-inhibition of chromatin regulators such as bromodomains, p300, MLL, HDACs, PRMT1, or DNMTs has been shown to synergize with EZH2 inhibitors across multiple cancer models by suppressing adaptive transcriptional programs and promoting pro-apoptotic gene expression^35,50,51^. These findings underscore the therapeutic potential of rational epigenetic combination strategies to overcome the intrinsic adaptability of cancer cells to PRC2 inhibition and highlight the need to identify context-specific dependencies that dictate response to EZH2-targeted therapies.

ATF4 is a central transcriptional effector of the integrated stress response, coordinating cellular adaptation to nutrient limitation, proteotoxic stress, and redox imbalance^52^. In cancer, ATF4 has been shown to support survival and therapy resistance by regulating amino acid metabolism, redox homeostasis, and stress-adaptive transcriptional programs^53^. Our data identify ATF4 as a key integrator of EZH2 inhibitor-induced stress in TNBC, linking epigenetic perturbation to metabolic reprogramming and altered amino acid utilization. While additional stress-responsive transcription factors may contribute to this response, our findings place ATF4-mediated metabolic adaptations at the center of the adaptive program that sustains cellular fitness under EZH2-inhibitory conditions. Notably, ATF4 has also been implicated in apoptotic signaling under conditions of severe or prolonged stress^54^, suggesting that the balance between adaptation and cell death following EZH2 inhibition is likely context-dependent and shaped by the magnitude and duration of ISR activation.

Metabolic rewiring is increasingly recognized as a consequence of EZH2 inhibition across cancer models^27,28^. Here, we demonstrate that in TNBC, EZH2 inhibition induces a stress-driven dependence on glutamine metabolism that can be therapeutically exploited. Given the limited specificity and clinical tractability of direct ISR or ATF4 inhibitors, we prioritized targeting glutaminase (GLS) as a downstream, druggable node of this adaptive metabolic program. Telaglenastat (CB-839), a clinically advanced GLS1 inhibitor, has shown activity in TNBC but with heterogeneous responses (#NCT02071862; NCT03057600)^55^, underscoring the need for predictive tumor contexts that define GLS dependency. Our findings indicate that EZH2 inhibition creates such a context, rendering TNBC cells and tumors selectively vulnerable to GLS inhibition. Whether additional EZH2-GLS combinations (eg, IACS-6274, BPTES) could refine efficacy remains to be investigated. Co-targeting EZH2 and glutamine metabolism, therefore, represents a rational strategy to enhance therapeutic efficacy, potentially reduce EZH2 inhibitor dosing, and broaden the clinical benefit of GLS-targeted therapies in TNBC.

Together, this study reframes EZH2 inhibition in TNBC as a stress-adaptive process rather than intrinsically cytotoxic therapy, revealing how cellular fitness is preserved through metabolic rewiring. By uncovering a targetable glutamine dependency that arises from this adaptation, our findings provide a conceptual and therapeutic framework for rational combination strategies that overcome stress tolerance and enhance the efficacy of epigenetic therapies in triple-negative breast cancer.

## KEY RESOURCES AND AVAILABILITY

Detailed information on reagents is provided in the Supplemental Materials and Methods. Further information and requests for reagents should be directed to the lead contact, Geneviève Deblois.

## EXPERIMENTAL MODEL AND STUDY PARTICIPANT DETAILS

### Cell culture

TNBC PDX-derived cells (GCRC1915 PDC) and the Hs 578T and MDA-MB-436 lines were maintained at 37°C in 5% CO₂. Hs 578T and MDA-MB-436 were cultured in Dulbecco’s Minimal Eagle’s Medium (DMEM) with 10% dialyzed FBS and 1% penicillin/streptomycin. GCRC1915 PDCs were cultured in Ham’s F12/DMEM (25:75) with 5% dialyzed FBS, 0.4 µg/mL hydrocortisone, 5 µg/mL insulin, 10 ng/mL EGF, 8.4 ng/mL cholera toxin, 50 µM ROCK inhibitor, and 50 µg/mL gentamycin. Cells were treated in vitro with UNC1999 (1 µM), EPZ-6438 (1 µM), Rapamycin (1 nM), Torin1 (1 nM), various doses of ISRIB, BSO, or CB-839, all dissolved in DMSO.

### Animal studies

All patient tissue samples were collected with informed consent under REB-approved protocols at McGill University Health Centre and Jewish General Hospital (SUR-2000-966; CER 2022-4027). Mice were maintained according to institutional animal care guidelines (SUR-99-780; 2014-7514). TNBC PDX tumors (GCRC1915 and GCRC1939) were cryopreserved in F12-DMEM with FBS and 10% DMSO at –80°C. Tumor fragments (∼1 mm³) were transplanted subcutaneously into the mammary fat pad of 5–7-week-old female non-obese diabetic scid gamma (NSG) mice. For cellular xenografts, 7.5 × 10⁵ Hs 578T cells were transplanted in 1:1 Matrigel. When tumors reached ∼150 mm³, mice were randomized to receive vehicle, UNC1999 (150–300 mg/kg, 5×/week), CB-839 (200 mg/kg, daily), or the combination. Tumor volume was measured using calipers [(length × width²)/2], and mice were sacrificed when tumors reached 2000–2500 mm³. UNC1999 and CB-839 were formulated in Sodium carboxymethyl cellulose (NaCMC) and (2-Hydroxypropyl)-β-cyclodextrin (HP-β-CD) with Tween 80, respectively.

## METHOD DETAILS

### RNA Extraction and Sequencing

PDX samples were collected after 21 days of treatment with or without UNC1999. Hs 578T and MDA-MB-436 cells were treated with EZH2 inhibitors (4 days) or siATF4 (2 days). RNA was extracted using Qiagen RNeasy kits, quantified with Qubit, and quality-checked on an Agilent Bioanalyzer. Poly-A-enriched or ribo-depleted RNA libraries were prepared with the KAPA RNA HyperPrep kit and sequenced on Illumina NextSeq500 (SE, 74–84 bp) or Novaseq (PE, 100 bp). Reads were trimmed with Trimmomatic and aligned to a hybrid human (GRCh38/Gencode37) and mouse (GRCm38/GencodeM25) genome using STAR. Human gene counts were normalized and analyzed for differential expression with DESeq2 (FDR < 0.05), and normalized counts were further used for functional enrichment analyses using GSEA with hallmark and GO biological pathways.

### Protein Extraction and Immunoblotting Analysis

Whole-cell pellets were lysed in buffer containing 50 mM Tris-HCl (pH 7.5), 150 mM NaCl, 1% Triton X-100, 0.1% SDS, and 0.5% sodium deoxycholate, sonicated (QSonica Q700, 4°C, 80% amplitude, 10 cycles of 30 s ON/30 s OFF), and quantified by Lowry assay. Proteins (20–40 µg) were separated on 8–12% SDS-PAGE gels and transferred to PVDF membranes (Trans-Blot Turbo, BioRad). Membranes were blocked with 5% milk or 3% BSA in TBS-T for 1 hour, incubated with primary antibodies overnight at 4°C, and detected using HRP-conjugated secondary antibodies and Clarity ECL substrate (BioRad).

### siRNA Transfection

Cells were seeded and transfected 4 hours later with siRNA (20 nM) using the siLentFect lipid reagent (BioRad) according to the manufacture’s protocol for a total duration of 48-72 h. 2 siRNAs/gene were used against *EIF2A2*/PKR, *EIF2A3*/PERK, *EIF2A/*eIF2α, *ATF4*, and non-Targeting siRNA (“si-Control” 1 and 2) were used as a negative control.

### Lentivirus preparation and generation of stable cell lines

MDA-MB-436 and Hs 578T EZH2 knockout cells were generated using lentiviral transduction. Cells were first transduced with lentiCas9-Blast and selected with blasticidin (10 µg/mL, 7 days). Two different sgRNAs targeting EZH2 exons 4 and 5, or a non-targeting control sgRNA (lenti-sgRNA puro), were then introduced and cells were selected with puromycin (1 µg/mL, 7 days). Lentiviral particles were produced in HEK 293T cells by co-transfecting psPAX2 and pVSV.G with the Cas9 or sgRNA vectors. EZH2 depletion was validated by immunoblotting.

### Chromatin Immunoprecipitation

Chromatin was prepared from UNC1999-treated PDX GCRC1915 tumors (300 mg/kg, 6×/week) and MDA-MB-436 cells (1 µM, 4 days) or vehicle controls. Cells and tissues were crosslinked with 1% formaldehyde (10 min), and chromatin was sonicated (QSonica Q700, 4°C, 80% amplitude, 35 cycles of 30 s ON/30 s OFF) to approximatively 400 bp fragments. Immunoprecipitation was performed with 50 µg chromatin and 4 µg antibody for H3K27me3/H3K27ac and 300 µg chromatin and 1 µg antibody for ATF4. Drosophila spike-in antibody (2 µg) was used with 50 µg of chromatin for histone mark normalization. IgG and input controls were included. ATF4 ChIP enrichment was verified by qPCR at *ASNS*, *SLC7A11*, *BCAT1*, and *PSAT1* loci (primer sequences listed in Supplemental Materials and Methods) using SensiFast SYBR. Purified DNA was quantified by Qubit and used to generate sequencing libraries (KAPA DNA HyperPrep Kit). Libraries were sequenced paired-end 100 bp, except one PDX replicate sequenced single-end 84 bp. ChIP-seq reads were aligned to human (GRCh38 Gencode40), mouse (GRCm39 GencodeM29), and Drosophila (BGDP.28) reference genomes to distinguish species-specific reads. Only uniquely mapped or properly paired reads were retained, and duplicates or low-quality reads (MAPQ < 30) were removed. Peaks were called using MACS3 with default FDR thresholds, using the --broad flag for H3K27me3. Spike-in normalization using Drosophila reads was applied to BigWig visualizations, and differential binding for H3K27ac samples was analyzed using DiffBind with DESeq2. Peak annotation and visualization were performed using ChIPseeker and deepTools.

### Transposable element expression

TE analysis included Salmon index generation, TE expression quantification and subfamily enrichment. TE sequences were extracted from RepeatMasker annotations, combined with transcriptomes, and indexed in Salmon for locus-specific quantification in human and mouse. TE annotations were refined using RepeatMasker, HMMER, and the Dfam database. Differential expression analysis was performed at the locus level using DESeq2, filtering for human TE insertions with sufficient read coverage. Subfamily enrichment was assessed by comparing differentially expressed TE insertions to the full annotated set using Fisher’s exact test. For full experimental and computational details, including all parameters, commands, and scripts, see the Supplemental Materials and Methods.

### Fixed cells immunofluorescence

Cells were plated on glass-bottom dishes, treated as indicated, fixed with methanol, permeabilized with Triton X-100, and blocked with PBS-T containing 3% BSA. Cells were incubated overnight at 4°C with anti-dsRNA antibody (1:100), followed by Alexa647-conjugated secondary antibody (1:500) and DAPI (300nM) nuclear staining. Images were acquired using a Leica SP8 confocal microscope and analyzed with ImageJ. dsRNA fluorescence intensity was quantified in 15 cells per biological replicate and normalized to DMSO-treated controls; IgG was used as a negative control.

### Tissue processing, immunohistochemistry and immunofluorescence and imaging

Four-micrometer sections were prepared from formalin-fixed, paraffin-embedded PDX tissues. Immunohistochemistry was performed using automated deparaffinization and antigen retrieval on a Discovery XT stainer, followed by primary antibody incubation using the Bond Polymer DAB Refine Kit on a Bond RX stainer and hematoxylin counterstaining. Slides were scanned using a NanoZoomer system and images were extracted with NDP Scan software. For immunofluorescence, sections were manually deparaffinized, rehydrated, and subjected to antigen retrieval in citrate buffer (pH 6.0). After blocking, sections were incubated overnight at 4°C with anti-dsRNA (1:50) and anti-H3K27me3 (1:100) antibodies, followed by Alexa488- and Alexa647-conjugated secondary antibodies (1:500) and DAPI (30nM) nuclear staining. Slides were imaged using a Cytation 5 system (20× objective). Image analysis was performed using QuPath, with cell identification based on DAPI intensity and nuclear morphology. Mean dsRNA signal intensity ratios were calculated and averaged across three biological replicates per condition.

### Proliferation assays

Cells were seeded in 96-well black plates (≥3 technical replicates) and incubated with AlamarBlue reagent (1:25, 3 h) on days 0, 3, and 6. Fluorescence was measured at 590 nm using a Spark® multimode plate reader, and relative growth was calculated as a ratio to day 0 for each condition.

### Viability assays

Cells were plated in 96-well plates (3 technical replicates) and treated as indicated. After treatment, cells were fixed with 100% cold methanol (10 min) and stained with Crystal Violet. Stained cells were lysed with 0.2% SDS, and absorbance was measured at 592 nm using a Spark® multimode plate reader.

### Protein misfolding assays

Misfolded and unfolded protein levels were measured using the tetraphenyl ethene maleimide (TPE-MI) probe, which fluoresces upon binding free cysteines. Cells were seeded in black 24-well plates and treated with or without EZH2 inhibitors for 48 h, then incubated with TPE-MI (25 µM) for 45 min. After PBS washes, fluorescence was measured at 470 nm using a Spark multimode plate reader and cells were imaged using a Cytation 5 system in brightfield and GFP channels.

### Extracellular metabolite consumption rate

Hs 578T and MDA-MB-436 cells were cultured in four independent biological replicates with or without UNC1999 or EPZ-6438. After 3 days, extracellular media were collected to measure glucose (g/L), lactate (g/L), glutamine (mmol/L) and glutamate (mmol/L) levels, and cells were counted for normalization. Media samples were clarified by centrifugation and analyzed using a BioProfile 400 Analyzer (Nova Biomedical). Metabolite concentrations were corrected by subtracting values from naïve media and normalized to cell number to calculate relative uptake or release under treated versus untreated conditions.

### Metabolomics and stable isotope tracing

For amino acid analysis, Hs 578T and MDA-MB-436 cells were cultured as described, treated with EZH2 inhibitors for 3 days, and transfected with siControl, siATF4-1, or siATF4-2. Extracellular media were extracted for metabolite analysis using ammonium formate washes and organic solvent extraction. For glutamine tracing, Hs 578T cells were treated with DMSO or UNC1999 for 2 days, then incubated in DMEM containing 4 mM 13C₅-glutamine or unlabeled glutamine for 2 h. Metabolites were extracted from cells as described above. Targeted metabolomics was performed by LC–MS using triple quadrupole MRM, while stable isotope tracer analyses were conducted using high-resolution QTOF mass spectrometry. Metabolites were separated using UHPLC with either ion-pairing reversed-phase chromatography for central carbon metabolites or hydrophilic interaction chromatography for amino acids. Data were normalized to total metabolite signal and analyzed for relative abundance and isotope enrichment. Detailed extraction procedures, chromatographic conditions, instrument parameters, and data analysis methods are provided in the Supplemental Materials and Methods.

### Extracellular measurement of oxygen consumption rate

Cells were treated with EZH2 inhibitors for 72 h prior to analysis. Oxygen consumption rate (OCR) and extracellular acidification rate (ECAR) were measured 3–4 times using a Seahorse XF-96 analyzer. Data are presented as OCR (pmol/min/µg protein) and ECAR (mpH/min/µg protein). Basal and maximal respiration were assessed under basal conditions following sequential injection of oligomycin (1 µM), FCCP (0.5 µM), and rotenone/antimycin A (0.5 µM) to determine ATP-linked respiration and maximal oxygen consumption.

## QUANTIFICATION AND STATISTICAL ANALYSIS

Data are presented as mean ± SD with no exclusions. Statistical analyses were performed using Prism 9 (GraphPad) and included two-tailed Student’s t-tests, one-way or two-way ANOVA with post hoc Dunnett’s tests, as indicated in figure legends. Analyses assumed normal distribution and equal variances. A P value ≤ 0.05 was considered significant. For in vivo studies, sample sizes were predetermined, and experiments were conducted with blinding.

## RESSOURCE AVAILABILITY

Genomic data is available under GSE307150 (elqhmuwyxjgxnsv), GSE307151 (wrcfumqslxgllcp), GSE307152 (ilczwwcivzynrqj), GSE307153 (shypoasyhpujzyv), GSE307154 (wngpkommttunhmh), GSE307155 (udolsiyappmxnev).

Code is available under https://github.com/annsophiegironne/ezh2_paper.

Requests for further information and resources should be directed to and will be fulfilled by the lead contact, Geneviève Deblois, Ph.D. Any additional information required to reanalyze the data reported in this paper is available from the lead contact upon request.

## AUTHOR CONTRIBUTION

Conceptualization: G.D., M.C., L.P.; Data curation: L.P., A.S.G., G.D.; Formal analyses: A.-S.G., L.P., M.C., G.D.; Funding acquisition: G.D., M.P.; Investigation: L.P., M.C., E.Q., G.A., H.P., Y.A., A.M., M.S.-A., M.D.T.R., F.G.,; Methodology: L.P., M.C., G.D., A.-S.G., S.L., D.A.; Resources: G.D., M.P., S.M.; Supervision: G.D.; Writing of original draft: G.D., M.C., L.P.; Review and editing: G.D., L.P., A.-S.G, S.L.

## ACKNOWLEDGEMENTS

We would like to thank all members of the Deblois lab for technical assistance and scientific input on the manuscript. The authors wish to thank the Institute for Research in Immunology and Cancer (IRIC) Genomic platform (Raphaëlle Lambert and Salwa Haidar) for ChIP-seq and RNA-seq libraries and sequencing, Histology platform (Marianne Isaac and Melina Narlis) for IHC staining. We thank the IRIC Bioinformatics platform for providing the infrastructure and assistance with the analyses. We acknowledge the Fonds de recherche du Québec (FRQ) – secteur Santé for supporting the Institute for Research in Immunology and Cancer (IRIC), an FRQ-designated research center. Metabolomics data were collected by the Metabolomics Innovation Resource (MIR) at the Rosalind and Morris Goodman Cancer Institute, McGill University. MIR is supported by our cost-recovery program, as well as the Terry Fox Research Institute, the Fondation du Cancer du sein du Québec, the Goodman Cancer Institute, and McGill University. (https://www.mcgill.ca/gci/facilities/metabolomics-innovation-resource-mir). This study was conducted with the support of the Canadian Institute for Health Research (CIHR) Funding Reference Number PJT-470501-CPT-CFCA-172840 (to G.D), the Réseau de Recherche en Cancer of the FRQ (to M. Park), Québec Breast Cancer Foundation (to M. Park) and the National Sciences and Engineering Research Councial of Canada (to S.M.). G. Deblois is a recipient of a CIHR Early Career Investigator in Cancer award, a Forbeck Scholar and a Research Scholar – Junior2 recipient from the Fonds de Recherche en Santé du Québec. M. Park is a James McGill Professor and holds the Diane and Sal Guerra Chair in Cancer Genetics at McGill University. L.P. M.F. and G.A. are recipients of a Fonds de Recherche en Santé du Québec doctoral scholarship. M.F. is a recipient of a CIHR doctoral studentship and a Canderel Doctoral Scholarship. A-S.G. and Y.A are recipients of a Fonds de Recherche en Santé du Québec Master’s scholarship, a CIHR BESC-M Master scholarship, and a Centennial scholarship from the Faculty of Pharmacy of the Université de Montréal.

## DECLARATION OF INTERESTS

The authors declare no competing interests.

---

**Supplementary Figure 1.**
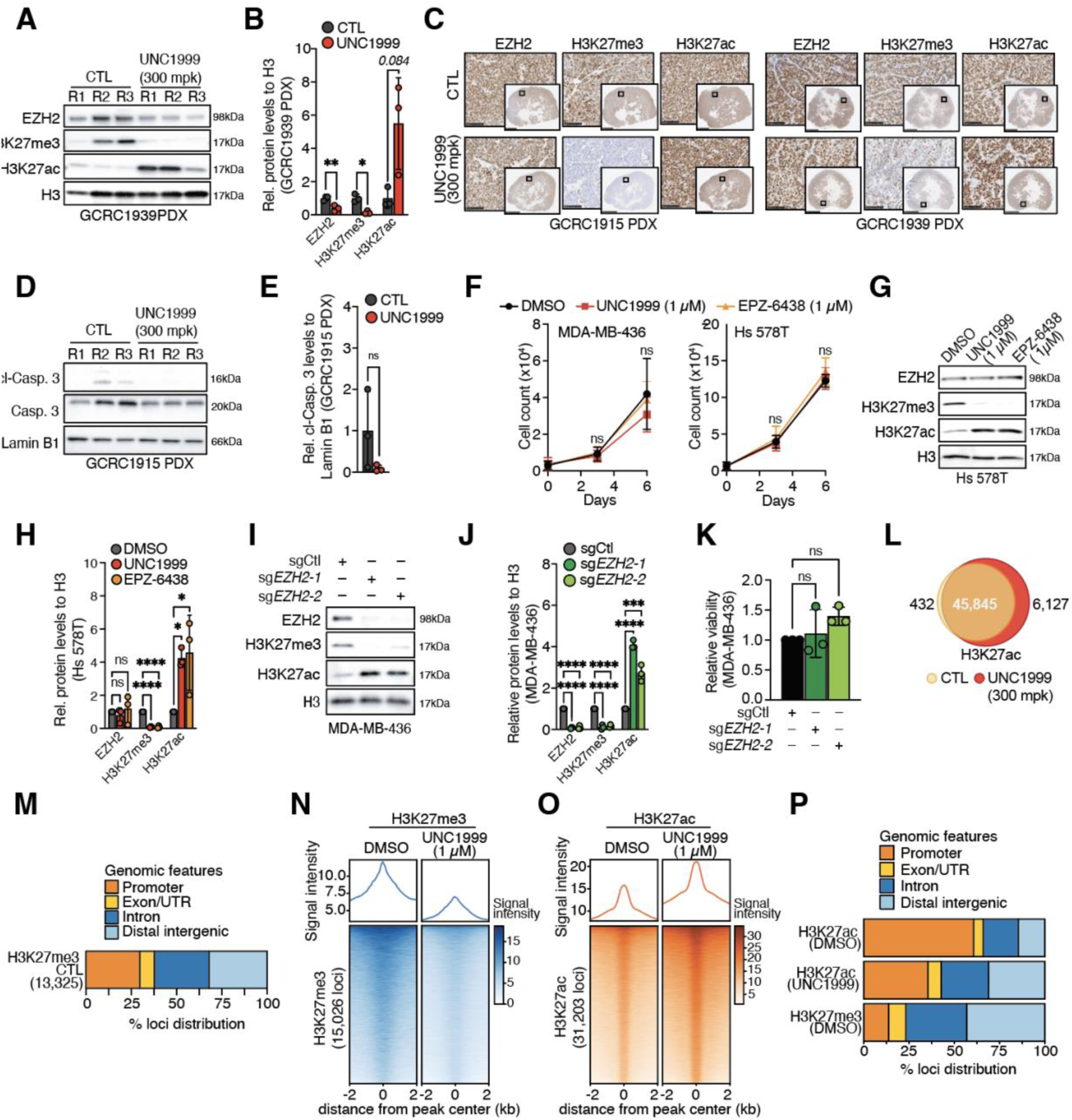
**A.** Immunoblot showing the levels of EZH2, H3K27me3, H3K27ac, and total H3 in GCRC1939 PDX exposed or not to UNC1999 (300 mg/kg; 21 days; N=3). **B.** Quantification of the relative protein levels from Fig.S1A (rel. to H3; N=3). **C.** Immunohistochemistry staining of EZH2, H3K27me3, and H3K27ac on GCRC1915 and GCRC1939 PDX exposed or not to UNC1999 (300 mg/kg, 21 days) (N=3); magnification, 20X; scale bars, 100µm. **D.** Immunoblot showing cl-Caspase 3, Caspase 3 and Lamin B1 levels in GCRC1915-PDX exposed or not to UNC1999 (300 mg/kg; 21 days) (N=3). **E.** Quantification of the relative cl-Caspase 3 levels from Fig.S1D (rel. to Lamin B1; N=3). **F.** Cell counts representing cell proliferation of MDA-MB-436 (left) and Hs 578T (right) cells exposed to UNC1999 or EPZ-6438 (1µM) or DMSO (0.2%) for up to 6 days (N=3). **G.** Immunoblot showing EZH2, H3K27me3, H3K27ac, and total H3 levels in Hs 578T cells exposed to UNC1999, EPZ-6438 (1 µM, 96 h; N=3) or mock treated (DMSO, 0.2%). **H.** Quantification of the relative protein levels from Fig.S1G (rel. to H3; N=3). **I.** Immunoblot showing EZH2, H3K27me3, H3K27ac, and total H3 levels in MDA-MB436-EZH2 knock-out cells using two sgRNAs targeting EZH2 or non-targeting sgRNA (sgCtl) (N=3). **J.** Quantification of the relative protein levels from Fig.S1I (rel. to H3; N=3). **K.** Relative viability (crystal violet) of MDA-MB-436 EZH2 knock-out cells using two different sgRNAs against *EZH2* (120h) or sgCtl (N=3). **L.** Venn diagrams representing the number of H3K27ac peaks that are common, specific to control, or to UNC1999 treatment (300 mg/kg, 21 days) in GCRC1915-PDX (N=2-3). **M.** Stacked bar charts illustrating the proportion of H3K27me3 peaks associated with distinct genomic localization/features relative to gene TSS upon control treatment in GCRC1915 PDX (300 mg/kg, 21 days; N=2-3). **N.** Heatmap of peak intensity signal of H3K27me3 ChIP-seq peaks (–2 to +2kbp window from peak center) in MDA-MB-436 cells treated with DMSO (0.2%; left) or UNC1999 (1µM; right) (N=3). Compiled tag density plots are illustrated at the top of each condition. **O.** Heatmap of peak intensity signal of H3K27ac upon DMSO (0.2%; left) or UNC1999 (1µM; right) treatment in MDA-MB-436 cells (N=2). Compiled tag density plots are illustrated at the top of each condition. **P.** Bar charts illustrating the proportion of H3K27me3 and H3K27ac peaks following DMSO (0.2%) or UNC1999 treatment (1µM; 96 h) in MDA-MB436 cells and associated with genomic features relative to gene TSS (N=2-3). Data represent the mean ± SD from 3 biological replicates (B, E, G, H, J and K). Statistical analysis was performed by two-tailed Student’s t test (B and E), one-way (G, J and K) and two-way ANOVA (H). ns, not significant, *p < 0.05, **p < 0.01, ***p < 0.001, ****p < 0.0001. R: replicate #.

**Supplementary Figure 2.**
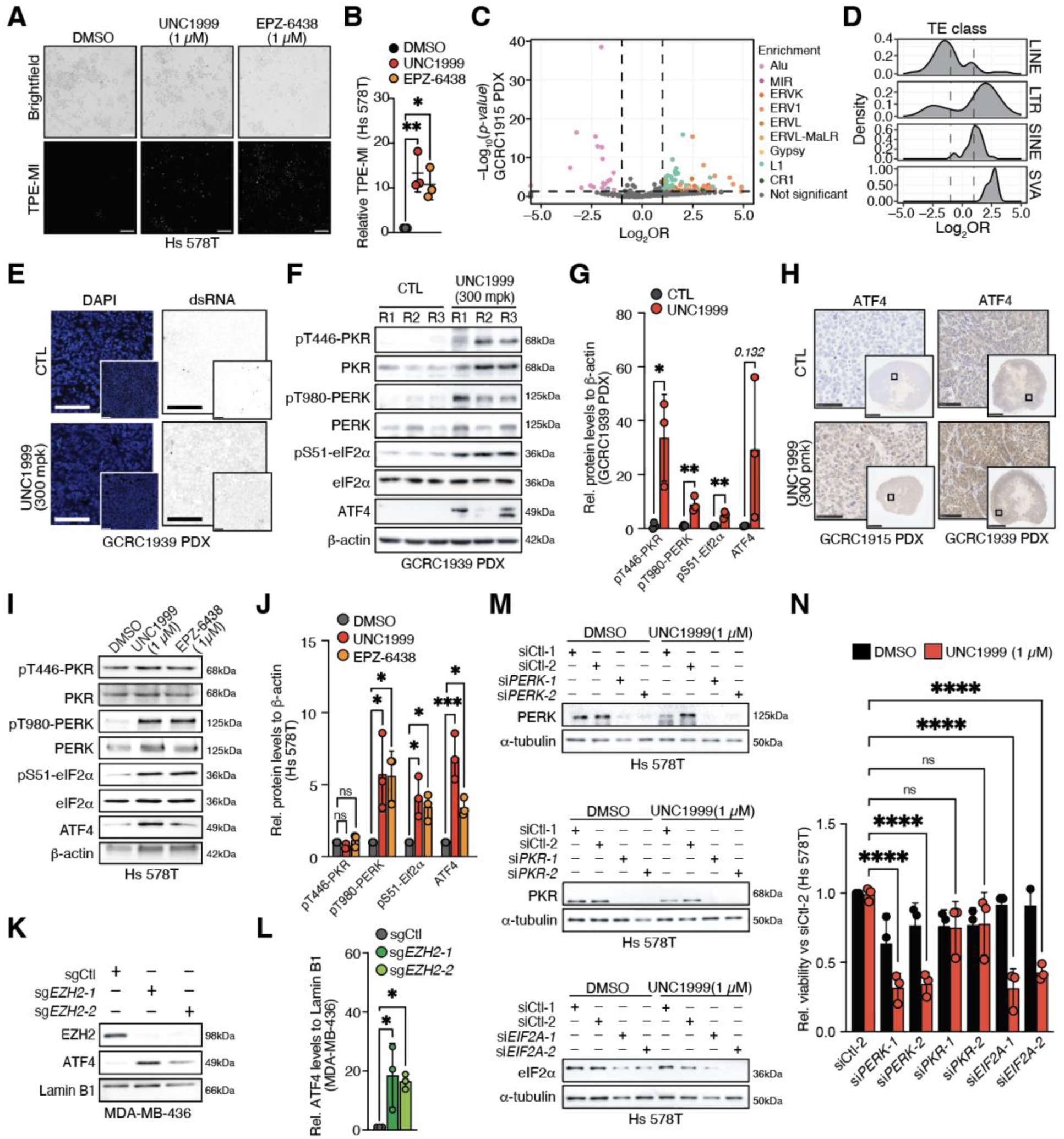
**A.** Detection of unfolded protein load in Hs 578T cells exposed to UNC1999 or EPZ-6438 (1µM; 48 h) or DMSO (0.2%; N=3), using the tetraphenyl ethene maleimide (TPE-MI) probe (IF) detecting misfolded protein loads. Cell density is shown in brightfield. Scale bar, 100µm. **B.** Quantification of misfolded protein load from Fig.S2A showing TPE-MI intensity normalized to naïve media and relative to DMSO (0.2%). **C.** Volcano plot showing enrichment level of significantly upregulated transposable elements (TE) insertions in UNC1999-treated GCRC1915 PDX (300 mg/kg; 21 days; N=3) relative to control. Colors correspond to their associated families. Odds ratio (OR) and p-value computed using Fisher’s exact test. **D.** Density chart representing global enrichment of TEs fully located within H3K27me3 peaks that gained H3K27ac following UNC1999 treatment in GCRC1915 PDX (N=2-3). Odds ratio (OR) and p-value computed using Fisher’s exact test. **E.** Immunofluorescence staining using J2 antibody in GCRC1939 PDX exposed to UNC1999 (300 mg/kg; 21 days) or control (N=3). Scale bar, 100µm. **F.** Immunoblot showing the levels of pT980-PERK, PERK, pT446-PKR, PKR, pS51-eIF2α, eIF2α, ATF4 and β-actin in GCRC1939-PDX exposed to UNC1999 (300 mg/kg; 21 days) or Control (N=3). **G.** Quantification of the relative protein levels from Fig. S2E (rel. to β-actin; N=3). **H.** IHC staining of ATF4 on GCRC1915 and GCRC1939 PDX exposed to UNC1999 (300 mg/kg, 21 weeks) or control (N=3). **I.** Immunoblot showing pT446-PKR, PKR, pT980-PERK, PERK, pS51-eIF2α, eIF2α, ATF4 and β-actin levels in Hs 578T exposed to control or UNC1999 (300 mg/kg; 21 days; N=3). **J.** Quantification of the relative protein levels from Fig.S2H (rel. to β-actin; N=3). **K.** Immunoblot showing EZH2, ATF4 and Lamin B1 levels in MDA-MB-436 EZH2 knock-out cells using two different sgRNAs (N=3). **L.** Quantification of the relative protein levels from Fig.S2J (rel. to Lamin B1; N=3). **M.** Immunoblot showing the levels of the indicated proteins (PKR, PERK, eIF2α, and α-tubulin) in Hs 578T cells exposed to UNC1999 (1µM; 48 h) or DMSO (0.2%; N=3) and depleted or not for the indicated gene/protein using two siRNAs/gene. **N.** Relative viability (crystal violet) of Hs 578T cells depleted of either PKR, PERK, or eIF2α using siRNAs (72 h) or siCtl and exposed to EZH2 inhibitor UNC1999 (1µM; 96 h) or DMSO (0.2%; N=3). Data shown are the mean ± SD from 3 biological replicates (B, G, J, L and N). Statistical analysis was performed by two-tailed Student’s t test (G), one-way (B, J and L) and two-way ANOVA (N). ns, not significant, *p < 0.05, **p < 0.01, ***p < 0.001, ****p < 0.0001. R: replicate #.

**Supplementary Figure 3.**
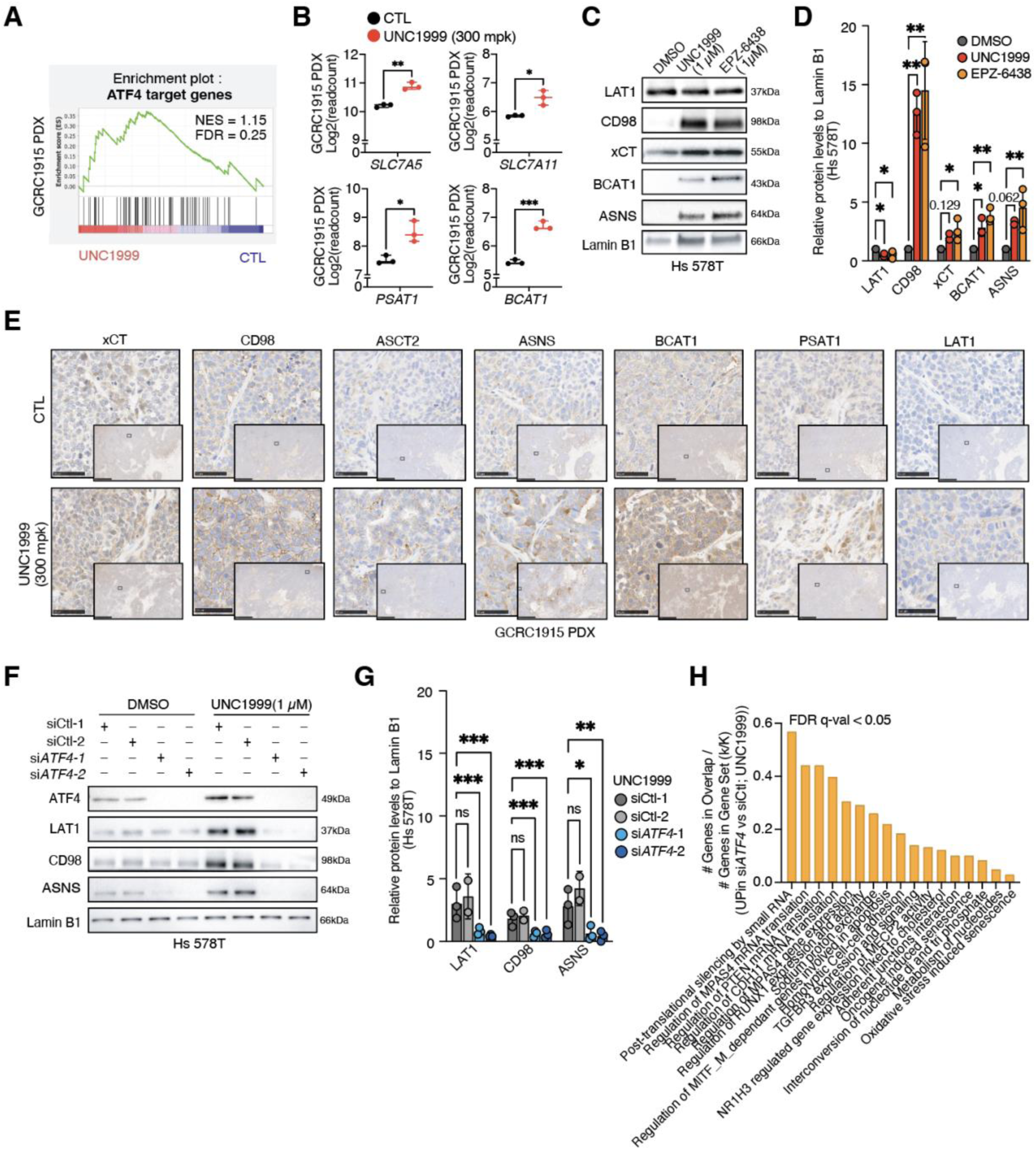
**A.** GSEA plot showing NES for *ATF4 targets* gene set generated from Koumenis et al.^60^ in GCRC1915 PDX exposed to UNC1999 (300 mg/kg, 21 days) or CTL (N=3). **B.** Gene expression levels of *SLC5A7*, *SLC7A11*, *PSAT1* and *BCAT1* in GCRC1915 PDX exposed to UNC1999 (300 mg/kg; 21 days) relative to control (N=3). **C.** Immunoblot showing LAT1, CD98, xCT, BCAT1, ASNS and Lamin B1 levels in Hs 578T exposed to DMSO (0.2%) or UNC1999 (1µM; 96 h; N=3). **D.** Quantification of the relative protein levels from Fig. S3C (rel. to Lamin B1; N=3). **E.** Immunohistochemistry staining of LAT1, CD98, xCT, ASCT2, PSAT1, BCAT1 and ASNS in GCRC1915 PDX exposed or not to UNC1999 (300 mg/kg; 21 days; N=3). Magnification, 2.5X and 40X. Scale bars, 50µm. **F.** Immunoblot showing the levels of ATF4, LAT1, CD98, ASNS and Lamin B1 in Hs 578T cells exposed to DMSO (0.2%) or UNC1999 (1µM; 72 h) (N=3) and depleted for ATF4 using 2 siRNAs (72 h). **G.** Quantification of the relative protein levels from Fig. S3F (rel. to Lamin B1; N=3). **H.** Bar graph representing down-regulated pathways (Reactome gene set) following ATF4 depletion in MDA-MB-436 cells exposed to UNC1999 (1µM; 48 h) relative to DMSO (0.2%; N=3). Data analysis is presented as the mean ± SD from three biological replicates (B, D and G). Statistical analysis was performed by two-tailed Student’s t test (B) and one-way ANOVA (D and G). ns, not significant, *p < 0.05 **p < 0.01, ***p < 0.001. R: replicate #.

**Supplementary Figure 4.**
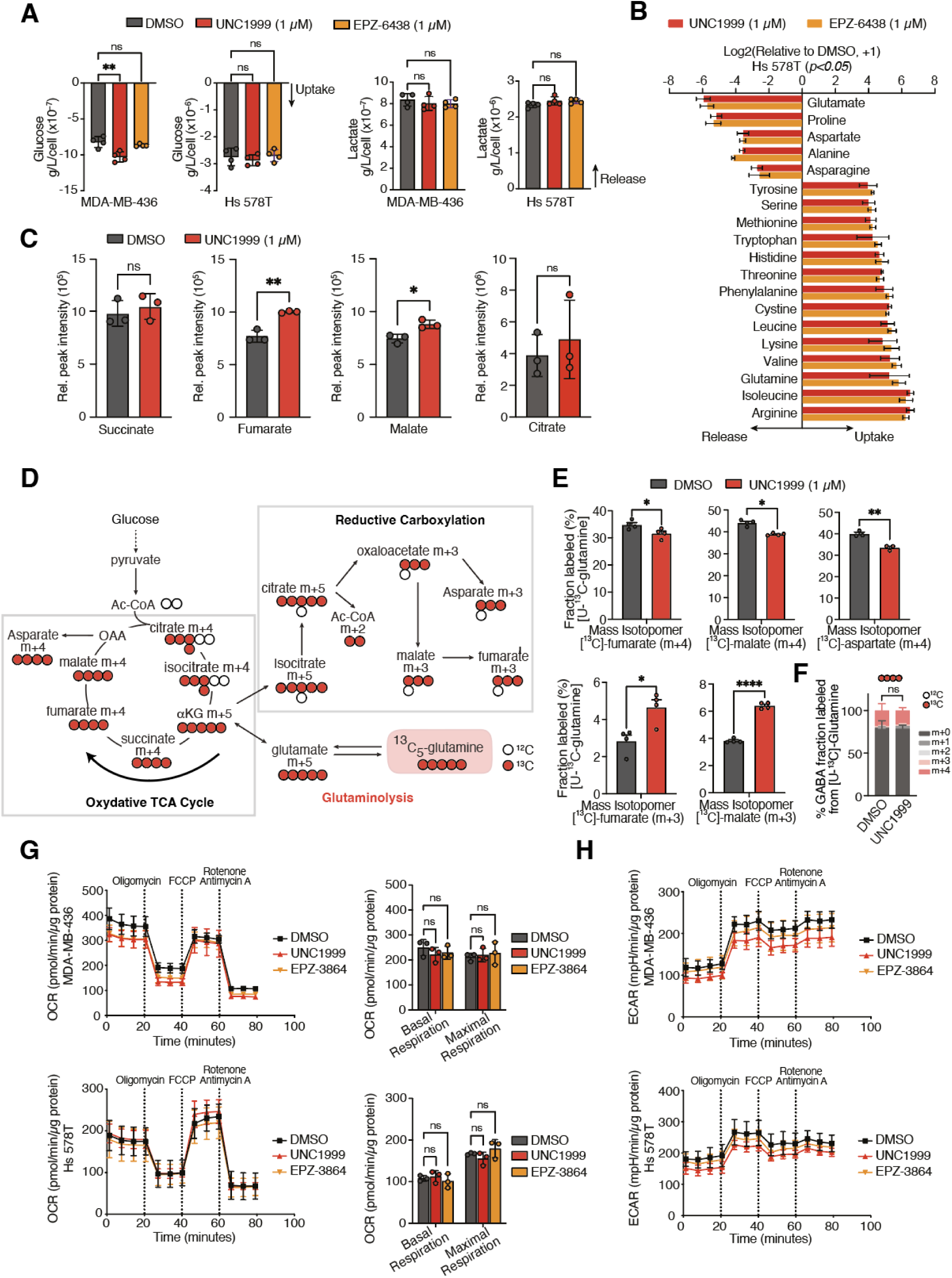
**A.** Relative quantification of glucose and lactate (BioProfile 400, Nova Biomedical) in conditioned media exposed to MDA-MB-436 or Hs 578T cells exposed to UNC1999 or EPZ-6438 (1µM; 72 h) or DMSO (0.2%; N=4; data shown relative to naïve media). **B.** Relative metabolite quantification in conditioned media using QQQ-LC-MS (Intrada-amino-acids) in Hs 578T cells exposed to UNC1999 and EPZ-6438 (1µM; 72 h) or DMSO (0.2%; N=4; p<0.05). Metabolite levels for each experimental condition are normalized to those in the naive medium. **C.** Quantification of specific intracellular metabolites in Hs 578T cells by QTOF-LC-MS (Ion-pairing) exposed to UNC1999 (1µM; 72 h) or DMSO (0.2%; N=3). **D.** Schematic representation of the ^13^C-labeling of glutaminolysis, oxidative TCA cycle, and reductive carboxylation intermediate from ^13^C_5_-glutamine. **E.** Percentage fraction of ^13^Carbon-labelled oxidative TCA cycle intermediates (top, fumarate m+4, malate m+4, and aspartate m+4), reductive carboxylation intermediates (bottom, malate m+3, fumarate m+3) in DMSO- and UNC1999-treated (1μM, 96 h) Hs 578T quantified by QTOF-LC-MS (ion pairing) after 2 h exposure to labelled ^13^C_5_-glutamine (N=4). **F.** Stack bar showing the fraction of ^13^C-labeled GABA isotopomers in Hs 578T cells exposed to UNC1999 (1μM, 96 h) or DMSO (0.2%; N=4). **G.** Left: Time-dependent Oxygen Consumption Rate (OCR) in MDA-MB-436 (top) and Hs 578T (bottom) cells exposed to UNC1999, EPZ-6438 (1µM; 72 h) or DMSO (0.2%; N=3) upon sequential addition of Oligomycin (1µM), Carbonyl cyanide-p-trifluoromethoxyphenylhydrazone (FCCP; 0.5µM) and Rotenone/AntimycinA (0.5µM). Right: Effect of treatment on OCR basal and maximal respiration is plotted. **H.** Time-dependent Extracellular Acidification Rate (ECAR) in MDA-MB-436 (top) and Hs 578T (bottom) cells exposed to UNC1999, EPZ-6438 (1µM; 72 h) or DMSO (0.2%; N=3) upon sequential addition of Oligomycin (1µM), FCCP (0.5µM) and Rotenone/AntimycinA (0.5µM). Data are the mean ± SD from 3 to 4 biological replicates (A, B, C, E, F, G and H). Statistical analysis was performed by two-tailed Student’s t test (C, E and F) and one-way ANOVA (A and G). ns, not significant, *p < 0.05, **p < 0.01, ****p < 0.0001.

**Supplementary Figure 5.**
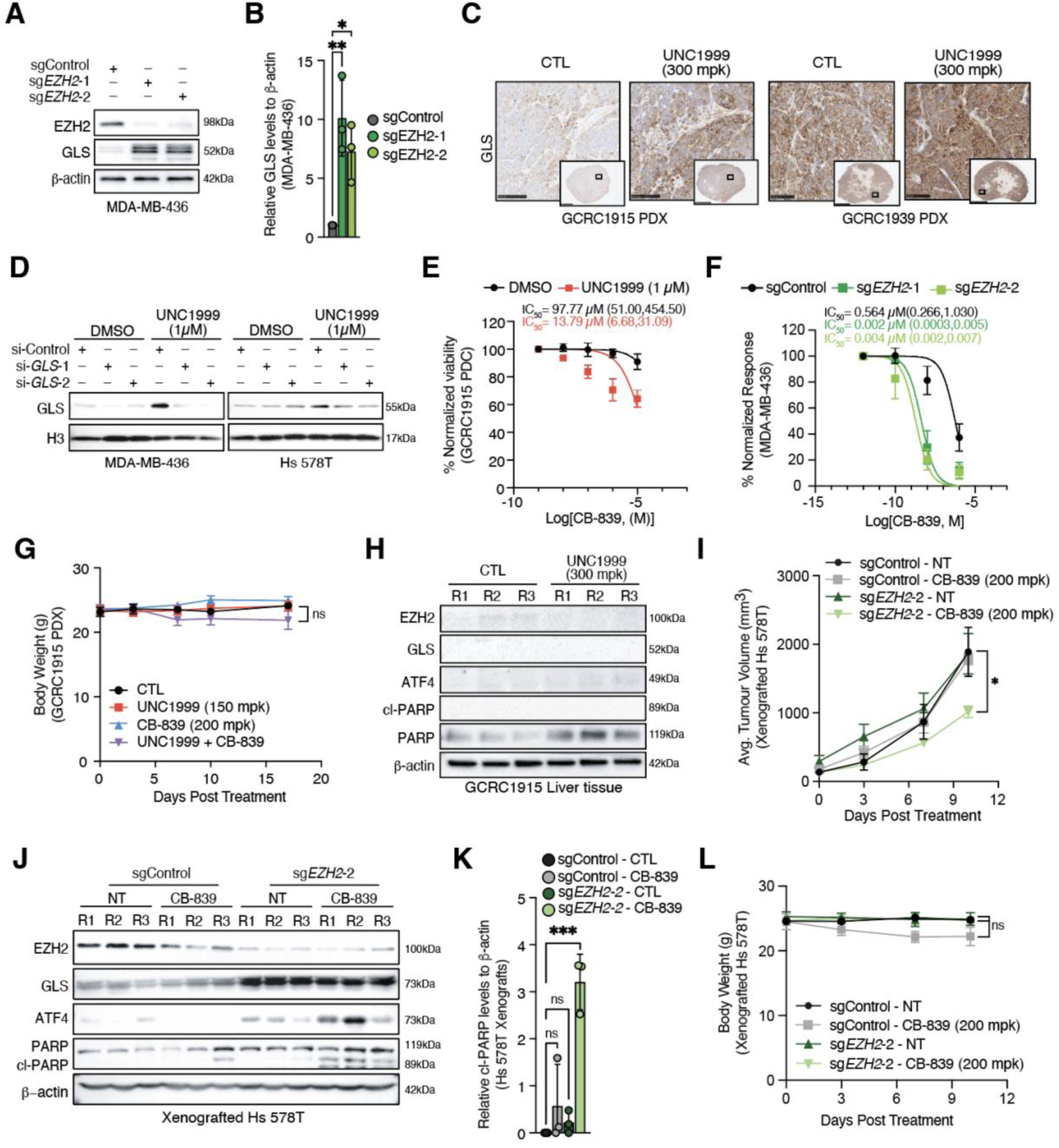
**A.** Immunoblot showing GLS and β-actin levels in MDA-MB-436 EZH2 knock-out cells using two different sgRNAs (N=3). **B.** Quantification of the relative GLS levels from Fig. S5A (rel. to β-actin; N=3). **C.** Immuno-histochemistry (IHC) staining for GLS in GCRC1915 and GCRC1939 PDXs exposed or not to UNC1999 (300 mg/kg, 21 days, N=3). Magnification, 20X; scale bars, 100µm. **D.** Immunoblot showing GLS and H3 levels in MDA-MB-436 or Hs 578T cells exposed to UNC1999 (1µM; 96 h) or DMSO (0.2%; N=3) and depleted of GLS using two siRNAs against *GLS* (96h) vs non-targeting siRNAs (siCtl). **E.** Dose-response curves representing the relative normalized viability of GCRC1915 PDC cells exposed to DMSO (0.2%) or UNC1999 (1µM; 120h) (N=3) and co-treated with increasing doses of CB-839. IC50 (95% Confidence Interval) was calculated and indicated at the top of the graph. **F.** Dose-response curves representing the relative normalized viability of MDA-MB-436 EZH2 knock-out cells and co-treated with increasing doses of CB-839 (120 h; N=3). IC50 (95% Confidence Interval) was calculated and indicated at the top of the graph. **G.** Body weight measurements of mice implanted with GCRC1915-PDX and treated as indicated (control, CB-839 (200 mg/kg; 21 days), UNC1999 (150 mg/kg; 21 days), or combination) over 10 days. **H.** Immunoblot showing the levels of the EZH2, GLS, ATF4, cl-PARP, and PARP of healthy liver from GCRC1939 PDX exposed to UNC1999 (300 mg/kg; 21 days) or control (N=3). **I.** Average tumor volume over time of sgControl and sg*EZH2* Hs 578T xenografted in female NSG mice exposed to CB-839 (200 mg/kg; 10 days) or control (N=3 per group). **J.** Immunoblot showing the levels of the EZH2, GLS, ATF4, cl-PARP, PARP, and β-actin in xenografted NSG mice either with sgControl or sg*EZH2* Hs 578T cells exposed to CB-839 (200 mg/kg; 10 days) or control (N=3). **O.** Quantification of the relative cl-PARP levels from Fig. S5I (rel. to β-actin; N=3). **L.** Body weight measurements of NGS mice xenografted with Hs 578T sgControl or sg*EZH2* and treated as indicated in I-J. The data shown are the mean ± SD from 3 biological replicates (B, E, F, G, K and L); error bars represent SEM for tumor volume (I). Statistical analysis was performed using one-way (B and K) and two-way ANOVA (G, I and L). ns, not significant, *p < 0.05, **p < 0.01, ***p < 0.001, ****p < 0.001. R: replicate #.

## Notes

The authors declare no potential conflicts of interest.

### Competing Interest Statement

The authors have declared no competing interest.

### Summary of Updates

Figure 1 revised to illustrate the induction of ISR Figure 4 revised to include mTOR activation

## REFERENCES

1. Su, Y. et al. Epigenetic reprogramming of epithelial mesenchymal transition in triple negative breast cancer cells with DNA methyltransferase and histone deacetylase inhibitors. J. Exp. Clin. Cancer Res. 37, 314 (2018).

2. Ratnam, N. M. et al. Reversing epigenetic gene silencing to overcome immune evasion in CNS malignancies. Front. Oncol. 11, 719091 (2021).

3. Verma, A. et al. EZH2-H3K27me3 mediated KRT14 upregulation promotes TNBC peritoneal metastasis. Nat. Commun. 13, 7344 (2022).

4. Pineda, B. et al. DNA methylation as an epigenetic signature predictive of response to neodjuvant treatment in TNBC patients. J. Clin. Oncol. 36, e12658–e12658 (2018).

5. Deblois, G. et al. Epigenetic Switch-Induced Viral Mimicry Evasion in Chemotherapy-Resistant Breast Cancer. Cancer Discov. 10, 1312–1329 (2020).

6. Marsolier, J. et al. H3K27me3 conditions chemotolerance in triple-negative breast cancer. Nat. Genet. 54, 459–468 (2022).

7. Comet, I., Riising, E. M., Leblanc, B. & Helin, K. Maintaining cell identity: PRC2-mediated regulation of transcription and cancer. Nat. Rev. Cancer 16, 803–810 (2016).

8. Lavarone, E., Barbieri, C. M. & Pasini, D. Dissecting the role of H3K27 acetylation and methylation in PRC2 mediated control of cellular identity. Nat. Commun. 10, 1679 (2019).

9. Chase, A. & Cross, N. C. P. Aberrations of EZH2 in cancer. Clin. Cancer Res. 17, 2613–2618 (2011).

10. Erokhin, M. et al. Clinical correlations of Polycomb repressive complex 2 in different tumor types. Cancers (Basel) 13, 3155 (2021).

11. Parreno, V. et al. Transient loss of Polycomb components induces an epigenetic cancer fate. Nature 629, 688–696 (2024).

12. Li, H., Cai, Q., Godwin, A. K. & Zhang, R. Enhancer of zeste homolog 2 promotes the proliferation and invasion of epithelial ovarian cancer cells. Mol. Cancer Res. 8, 1610–1618 (2010).

13. Bachmann, I. M. et al. EZH2 expression is associated with high proliferation rate and aggressive tumor subgroups in cutaneous melanoma and cancers of the endometrium, prostate, and breast. J. Clin. Oncol. 24, 268–273 (2006).

14. Morin, R. D. et al. Somatic mutations altering EZH2 (Tyr641) in follicular and diffuse large B-cell lymphomas of germinal-center origin. Nat. Genet. 42, 181–185 (2010).

15. Gu, Z. et al. Loss of EZH2 Reprograms BCAA Metabolism to Drive Leukemic Transformation. Cancer Discov. 9, 1228–1247 (2019).

16. Ernst, T. et al. Inactivating mutations of the histone methyltransferase gene EZH2 in myeloid disorders. Nat. Genet. 42, 722–726 (2010).

17. Zeng, D., Liu, M. & Pan, J. Blocking EZH2 methylation transferase activity by GSK126 decreases stem cell-like myeloma cells. Oncotarget 8, 3396–3411 (2017).

18. Daisuke, S. et al. EZH2 inhibitor DZNep induces apoptosis in adult T-cell leukemia/lymphoma cells by BCL2 suppression via regulation of Mir-181a. Blood 122, 4265–4265 (2013).

19. Caganova, M. et al. Germinal center dysregulation by histone methyltransferase EZH2 promotes lymphomagenesis. J. Clin. Invest. 123, 5009–5022 (2013).

20. Keller, P. J. et al. Comprehensive target engagement by the EZH2 inhibitor tulmimetostat allows for targeting of ARID1A mutant cancers. Cancer Res. 84, 2501–2517 (2024).

21. Huang, R., Wu, Y. & Zou, Z. Combining EZH2 inhibitors with other therapies for solid tumors: more choices for better effects. Epigenomics 14, 1449–1464 (2022).

22. Hoy, S. M. Tazemetostat: First Approval. Drugs 80, 513–521 (2020).

23. Julia, E. & Salles, G. EZH2 inhibition by tazemetostat: mechanisms of action, safety and efficacy in relapsed/refractory follicular lymphoma. Future Oncol. 17, 2127–2140 (2021).

24. Piunti, A. et al. Immune activation is essential for the antitumor activity of EZH2 inhibition in urothelial carcinoma. Sci. Adv. 8, eabo8043 (2022).

25. Chibaya, L. et al. EZH2 inhibition remodels the inflammatory senescence-associated secretory phenotype to potentiate pancreatic cancer immune surveillance. *Nat*. Cancer 4, 872–892 (2023).

26. Bowen, C. M., et al. Inhibition of histone methyltransferase EZH2 for immune interception of colorectal cancer in Lynch syndrome. JCI Insight 10, (2025).

27. Nylund, P. et al. A distinct metabolic response characterizes sensitivity to EZH2 inhibition in multiple myeloma. Cell Death Dis. 12, 167 (2021).

28. Nocito, M. C. et al. A targetable antioxidant defense mechanism to EZH2 inhibitors enhances tumor cell vulnerability to ferroptosis. Cell Death Dis. 16, 291 (2025).

29. W-M Fan, T., et al. Metabolic reprogramming driven by EZH2 inhibition depends on cell-matrix interactions. J. Biol. Chem. 300, 105485 (2024).

30. Xu, L. et al. Pharmacological inhibition of EZH2 combined with DNA-damaging agents interferes with the DNA damage response in MM cells. Mol. Med. Rep. 19, 4249–4255 (2019).

31. Bianchini, G., Balko, J. M., Mayer, I. A., Sanders, M. E. & Gianni, L. Triple-negative breast cancer: challenges and opportunities of a heterogeneous disease. Nat. Rev. Clin. Oncol. 13, 674–690 (2016).

32. Yomtoubian, S. et al. Inhibition of EZH2 Catalytic Activity Selectively Targets a Metastatic Subpopulation in Triple-Negative Breast Cancer. Cell Rep. 30, 755–770.e6 (2020).

33. De Brot, M., Rocha, R. M., Soares, F. A. & Gobbi, H. Prognostic impact of the cancer stem cell related markers ALDH1 and EZH2 in triple negative and basal-like breast cancers. Pathology 44, 303–312 (2012).

34. Yu, Y. et al. Epigenetic Co-Deregulation of EZH2/TET1 is a Senescence-Countering, Actionable Vulnerability in Triple-Negative Breast Cancer. Theranostics 9, 761–777 (2019).

35. Huang, X. et al. Targeting Epigenetic Crosstalk as a Therapeutic Strategy for EZH2-Aberrant Solid Tumors. Cell 175, 186–199.e19 (2018).

36. Yamagishi, M. et al. Mechanisms of action and resistance in histone methylation-targeted therapy. Nature 627, 221–228 (2024).

37. Bai, Y. et al. Inhibition of enhancer of zeste homolog 2 (EZH2) overcomes enzalutamide resistance in castration-resistant prostate cancer. J. Biol. Chem. 294, 9911–9923 (2019).

38. Luo, C. et al. H3K27me3-mediated PGC1α gene silencing promotes melanoma invasion through WNT5A and YAP. J. Clin. Invest. 130, 853–862 (2020).

39. Deb, G., Thakur, V. S. & Gupta, S. Multifaceted role of EZH2 in breast and prostate tumorigenesis: epigenetics and beyond. Epigenetics 8, 464–476 (2013).

40. Mahara, S. et al. HIFI-α activation underlies a functional switch in the paradoxical role of Ezh2/PRC2 in breast cancer. Proc. Natl. Acad. Sci. U. S. A. 113, E3735–44 (2016).

41. da Hora, C. C. et al. Sustained NF-κB-STAT3 signaling promotes resistance to Smac mimetics in Glioma stem-like cells but creates a vulnerability to EZH2 inhibition. Cell Death Discov. 5, 72 (2019).

42. Dann, S. G. et al. Reciprocal regulation of amino acid import and epigenetic state through Lat1 and EZH2. EMBO J. 34, 1773–1785 (2015).

43. Anwar, T., Gonzalez, M. E. & Kleer, C. G. Noncanonical Functions of the Polycomb Group Protein EZH2 in Breast Cancer. Am. J. Pathol. (2021) doi:10.1016/j.ajpath.2021.01.013.

44. Knutson, S. K. et al. A selective inhibitor of EZH2 blocks H3K27 methylation and kills mutant lymphoma cells. Nat. Chem. Biol. 8, 890–896 (2012).

45. Hsieh, Y.-Y., Lo, H.-L. & Yang, P.-M. EZH2 inhibitors transcriptionally upregulate cytotoxic autophagy and cytoprotective unfolded protein response in human colorectal cancer cells. Am. J. Cancer Res. 6, 1661–1680 (2016).

46. Morel, K. L. et al. EZH2 inhibition activates a dsRNA-STING-interferon stress axis that potentiates response to PD-1 checkpoint blockade in prostate cancer. *Nat*. Cancer 2, 444–456 (2021).

47. Qiu, F. et al. EZH2 inhibition activates dsRNA-interferon axis stress and promotes response to PD-1 checkpoint blockade in NSCLC. J. Cancer 13, 2893–2904 (2022).

48. Chomiak, A. A. et al. Select EZH2 inhibitors enhance viral mimicry effects of DNMT inhibition through a mechanism involving NFAT:AP-1 signaling. Sci. Adv. 10, eadk4423 (2024).

49. Kazansky, Y. et al. Overcoming clinical resistance to EZH2 inhibition using rational epigenetic combination therapy. Cancer Discov. 14, 965–981 (2024).

50. Li, Z. et al. Methylation of EZH2 by PRMT1 regulates its stability and promotes breast cancer metastasis. Cell Death Differ. 27, 3226–3242 (2020).

51. Zhang, L. et al. DNMT and EZH2 inhibitors synergize to activate therapeutic targets in hepatocellular carcinoma. Cancer Lett. 548, 215899 (2022).

52. Lines, C. L., McGrath, M. J., Dorwart, T. & Conn, C. S. The integrated stress response in cancer progression: a force for plasticity and resistance. Front. Oncol. 13, 1206561 (2023).

53. Wortel, I. M. N., van der Meer, L. T., Kilberg, M. S. & van Leeuwen, F. N. Surviving stress: Modulation of ATF4-mediated stress responses in normal and malignant cells. Trends Endocrinol. Metab. 28, 794–806 (2017).

54. Matsumoto, H. et al. Selection of autophagy or apoptosis in cells exposed to ER-stress depends on ATF4 expression pattern with or without CHOP expression. Biol. Open 2, 1084–1090 (2013).

55. Harding, J. J. et al. A phase I dose-escalation and expansion study of telaglenastat in patients with advanced or metastatic solid tumors. Clinical cancer research: an official journal of the American Association for Cancer Research vol. 27 4994–5003 (2021).

56. Tameire, F. et al. ATF4 couples MYC-dependent translational activity to bioenergetic demands during tumour progression. Nat. Cell Biol. 21, 889–899 (2019).

