## Supplementary Material and Methods for "EZH2 Inhibition Induces a Metabolic Stress Response Sensitizing TNBC to Glutaminase Targeting"

[7] Lead contact.

* To whom correspondence should be addressed.

### SUPPLEMENTARY RESSOURCES TABLE

| REAGENT or RESOURCE | SOURCE | IDENTIFIER |
|---|---|---|
| **Antibodies** | | |
| EZH2 | Cell Signaling Technology | Cat#5246; RRID: AB_10694683 |
| H3K27me3 | Cell Signaling Technology | Cat#9733; RRID: AB_2616029 |
| H3K27Ac | Abcam | Cat#ab4729; RRID: AB_2118291 |
| H3 | Abcam | Cat#ab10799; RRID: AB_470239 |
| dsRNA | EMD Millipore | Cat#MABE-1134; RRID: AB_2819101 |
| pT446-PKR | Abcam | Cat#ab32036; RRID: AB_777310 |
| PKR | Cell Signaling Technology | Cat#12297; RRID: AB_2665515 |
| pT980-PERK | Cell Signaling Technology | Cat#3179; RRID: AB_2095853 |
| PERK | Cell Signaling Technology | Cat#3192; RRID: AB_2095847 |
| β-actin | Sigma-Aldrich | Cat#A5441; RRID: AB_476744 |
| pS51-eIF2α | Cell Signaling Technology | Cat#9721; RRID: AB_330951 |
| eIF2α | Cell Signaling Technology | Cat#9722; RRID: AB_2230924 |
| ATF4 (IHC) | Abcam | Cat#ab31390; RRID: AB_725568 |
| ATF4 (immunoblot and ChIP) | Cell Signaling Technology | Cat#11815; RRID: AB_2616025 |
| α-tubulin | Sigma-Aldrich | Cat#T5168; RRID: AB_477579 |
| Cleaved-PARP | Cell Signaling Technology | Cat#9541; RRID: AB_331426 |
| PARP | Cell Signaling Technology | Cat#9542; RRID: AB_2160739 |
| Lamin B1 | Abcam | Cat#ab16048; RRID: AB_443298 |
| LAT1 | Cell Signaling Technology | Cat#32683; RRID: AB_3265513 |
| CD98 | Cell Signaling Technology | Cat#47213; RRID: AB_2799323 |
| ASNS | Cell Signaling Technology | Cat#92479; RRID: AB_3082991 |
| BCAT1 (for immunoblot) | Cell Signaling Technology | Cat#12822; RRID: AB_2798035 |
| BCAT1 (for IHC) | Origene | Cat#TA504360; RRID: AB_11126287 |
| PSAT1 | NOVUS Biological | Cat#NBP1-32920; RRID: AB_2172600 |
| GLS | Abcam | Cat#ab156876; RRID: AB_2721038 |
| pS1859-CAD | Cell Signaling Technology | Cat#67235; RRID: AB_2799722 |
| CAD | Cell Signaling Technology | Cat#11933; RRID: AB_2797772 |
| pS235/236-S6 | Cell Signaling Technology | Cat#2211; RRID: AB_331679 |
| S6 | Santa Cruz Biotechnology | Cat#sc-74459; RRID: AB_1129205 |
| xCT | Cell Signaling Technology | Cat#NB300-318SS; RRID: AB_921456 |
| IgG anti-mouse HRP | Cell signaling Technology | Cat#7076; RRID: AB_330924 |
| IgG anti-rabbit HRP | Cell signaling Technology | Cat#; 7074; RRID: AB_2099233 |
| IgG anti-mouse (Alexa 647) | Abcam | Cat#ab150119; RRID: AB_2811129 |
| IgG anti-rabbit (Alexa 488) | Abcam | Cat#ab150081; RRID: AB_2734747 |
| IgG Rabbit isotype control | Cell Signaling Technology | Cat#3900; RRID: AB_1550038 |
| IgG Mouse isotype control | Cell Signaling Technology | Cat#5415; RRID: AB_10829607 |
| **Bacterial strain** | | |
| TOP10 competent *E. coli* | Thermo Fisher Scientific | Cat#C404003 |
| **Chemicals, peptides, and recombinant proteins** | | |
| UNC1999 | UHN Shanghai | Cat#1431612-23-5 |
| EPZ-6438 | MedChem Express | Cat#HY-13803 |

| | | |
|---|---|---|
| ISRIB | EMD Millipore | Cat#509584 |
| BSO | EMD Millipore | Cat#B2515 |
| CB-839 | EMD Millipore | Cat#5.33717.0001 |
| Rapamycin | MedChem Express | Cat#HY-10219 |
| Torin1 | MedChem Express | Cat#HY-13003 |
| Blasticidine | Wisent | Cat#400-190-EM |
| Puromycin | Wisent | Cat#400-160-EM |
| Cristal Violet | Sigma Aldrich | Cat#C0765 |
| Alamar Blue | Thermo Fisher Scientific | Cat#DAL1025 |
| Formaldehyde | Sigma Aldrich | Cat#252549 |
| Dynabeads A | Thermo Fisher Scientific | Cat#10004D |
| Dynabeads G | Thermo Fisher Scientific | Cat#10002D |
| Spike-in antibody | Active Motif | Cat#61686 |
| Spike-in Chromatin | Active Motif | Cat#53083 |
| DAPI | Thermo Fisher Scientific | Cat#D1306 |
| TPE-MI | MedChem Express | Cat#HY-143218 |
| SensiFast SYBR | Bioline | Cat#BIO-98020 |
| siLenFect lipid reagent | Bio-rad | Cat#1703361 |
| L-Glutamine-$^{13}C_5$ | Sigma Aldrich | Cat#605166, CAS: 184161-19-1 |
| Matrigel | Corning | Cat#354248 |
| DMEM | Gibco | Cat#11995-065 |
| Ham's F-12 | Gibco | Cat#11765-054 |
| Penicillin–Streptomycin | Wisent | Cat#098-150 |
| FBS dialyzed | Wisent | Cat#080-950 |
| EGF | BPS Bioscience | Cat# 90201-1 |
| Insulin | Gibco | Cat#12585014 |
| Hydrocortisone | Multicell | Cat#511-012 |
| Rock inhibitor | Enzo Life Sciences | Cat#Y-27632 |
| Cholera toxin | Sigma Aldrich | Cat#C8052 |
| Gentamycin | Gibco | Cat#**15-710-072** |
| **Kit** | | |
| Perm/Fix | BD Biosciences | Cat#554723 |
| Seahorse FluxPaks | Agilent | Cat#103022-100 |
| Monarch Genomic DNA purification kit | New England BioLabs | Cat#T3010S |
| RNeasy Fibrous Tissue Mini Kit | Qiagen | Cat#74704 |
| RNeasy Mini Kit | Qiagen | Cat#74104 |
| SensiFast SYBR No-ROX kit | FroggaBio | Cat#BIO-98020 |
| **Experimental models: Cell lines and models** | | |
| Human: Hs 578T | ATCC | *RRID:CVCL_0332* |
| Human: MDA-MB-436 | ATCC | *RRID:CVCL_0623* |
| Human: MDA-MB-436 sgControl | This study | N/A |
| Human: MDA-MB-436 sgEZH2-1 | This study | N/A |

| | | |
|---|---|---|
| Human: MDA-MB-436 sgEZH2-1 | This study | N/A |
| Human: Hs 578T sgControl | This study | N/A |
| Human: Hs 578T sgEZH2-2 | This study | N/A |
| Human: HEK293T | ATCC | RRID:CVCL_0063 |
| Human: PDC1915 (derived from GCRC1915 human tumor) | Savage et al.[59] | N/A |
| Human: GCRC1915 | Savage et al.[59] | N/A |
| Human: GCRC1939 | Savage et al.[59] | N/A |
| Mouse: NOD.Cg-PrkdcscidIl2rgtm1Wjl/SzJ | The Jackson Laboratory | Cat#JAX:005557 |
| **Recombinant DNA and RNA material** | | |
| lentiCas9-Blast (3rd generation lentiviral production) | Addgene | Plasmid #52962 |
| psPAX2 (2nd generation lentiviral production) | Addgene | Plasmid #12260 |
| pVSV.G (2nd generation lentiviral production) | Addgene | Plasmid #8454 |
| sgEZH2-1 (5'-GTTGGAAAATCCAAGTCACTGG-3') | Sigma aldrich | Sanger Arrayed Whole Genome Lentiviral CRISPR Library (ID: HS5000005721) |
| sgEZH2-2 (5'-GGAGATGAAGTTTTAGATCAG-3) | Sigma aldrich | Sanger Arrayed Whole Genome Lentiviral CRISPR Library (ID: HS5000005722) |
| Lenti-sgRNA puro (Non targeting sgRNA) | Addgene | Plasmid #104990 |
| si-EIF2AK3/PERK-1 | Thermo Fisher Scientific | Cat#4392420, ID: s18101 |
| si-EIF2AK3/PERK-2 | Thermo Fisher Scientific | Cat#4392420, ID: s18103 |
| si-EIF2AK2/PKR-1 | Thermo Fisher Scientific | Cat#4392420, ID: s229500 |
| si-EIF2AK2/PKR-2 | Thermo Fisher Scientific | Cat#4392420, ID: s229501 |
| si-EIF2A/eIF2α-1 | Thermo Fisher Scientific | Cat#4392420, ID: s38345 |
| si-EIF2A/eIF2α-2 | Thermo Fisher Scientific | Cat#4392420, ID: s38346 |
| si-ATF4-1 | Thermo Fisher Scientific | Cat#4392420, ID: s1702 |
| si-ATF4-2 | Thermo Fisher Scientific | Cat#4392420, ID: s1703 |
| si-GLS-1 | Thermo Fisher Scientific | Cat#4392420, ID: s5838 |
| si-GLS-2 | Thermo Fisher Scientific | Cat#4392420, ID: s5839 |
| **Software and algorithms** | | |
| FlowJo (v10.10) | Treestar | https://www.flowjo.com/ |
| ImageJ (v2.14) | NIH | https://imagej.net/software/fiji/ |
| Prism (v9.1.1) | GraphPad | https://www.graphpad.com/ |
| MACS (v3.0) | Github | https://github.com/macs3-project/MACS |
| Trimmomatic (v0.39) | Usadel Lab | http://www.usadellab.org/cms/?page=trimmomatic |
| bwa (v0.7.17) | Github | https://github.com/lh3/bwa |
| QuPath (v0.5.0) | Github | https://github.com/qupath/qupath/releases |

#### SUPPLEMENTARY METHOD DETAILS

##### Salmon Index Generation

To build the custom Salmon index used for quantification, transposable element (TE) sequences were extracted based on the RepeatMasker annotations. A BED file containing the genomic coordinates of TE loci was generated, and the corresponding sequences were retrieved from the human and mouse reference genomes using BEDTools getfasta (v2.30.0).

These TE sequences were then combined with transcriptome FASTA files from both human and mouse annotations. The final index was built using Salmon (v1.10.2) with a k-mer size of 31, keeping duplicate sequences and without clipping polyA tails. This combined index allowed accurate quantification of both species-specific transcripts and locus-specific TE insertions, and was designed to facilitate discrimination between human and mouse reads in PDX RNA-seq data.

##### Detection of TE Expression

Transposable element (TE) annotations were generated using RepeatMasker (v4.1.5). Annotation was performed by combining RepeatMasker with the HMMER tool (v3.3.1) and the Dfam database (v3.7). The output file from RepeatMasker in fa.out.gff format was converted to a GTF file using a script provided by the Hammell lab (https://github.com/mhammell-laboratory). Annotation was performed using the following command: perl RepeatMasker -species [human/mouse] -gff -pa 32 -u -xm [fasta_genome_file] -dir [output_dir].

For RNA-seq data from PDX samples, human and mouse transcript annotations were merged with locus-specific human and mouse TE annotations, in order to distinguish the origin of sequencing reads (human vs. mouse, transcript vs TE). Quantification was performed at the locus level using Salmon (v1.10.2). Differential expression analysis was performed exclusively for human TE insertions using DESeq2 (v1.42.0), using a threshold of log2FoldChange > 1 and an adjusted p-value < 0.05. Insertions were filtered for those with at least three samples having a minimum of 10 reads.

##### TE Subfamily Enrichment Analysis

Enrichment of TE subfamilies was assessed by comparing the proportion of differentially upregulated human TE insertions across subfamilies to their proportion in the full set of annotated and analyzed TE loci. For each subfamily, insertions were classified into four categories based on two criteria: (i) whether the insertion was differentially expressed or not, and (ii) whether it belonged to the subfamily of interest or not. This comparison was performed using a two-sided Fisher's exact test implemented in R (v4.3.3), considering the total set of TE insertions annotated by RepeatMasker and included in the analysis as the background. Subfamilies were considered significantly enriched if they showed an odds ratio (OR) $\geq 2$ and a p-value $< 0.05$.

**Metabolite extraction for LC-MS**. Extracellular medium was collected for differential amino acid analysis, and in the case of tracing, cells were washed with 150 mM (pH=7.4) ammonium formate buffer containing N-ethylmaleimide (NEM) to protect thiols, harvested with 50% MeOH/50% LC/MS-grade water (with NEM). To both sample types, acetonitrile (ACN) was added and cells homogenized by bead beating. Lastly, the samples were partitioned using dichloromethane and water. Polar (aqueous layer) metabolites were dried in a sample cooled (-4°C) vacuum concentrator overnight. The samples were stored at −80°C until further analysis was performed.

**LC-MS.** For the detection of central carbon metabolites two chromatographic methods were employed using Agilent 6470 Triple Quadrupole (QQQ) instruments for targeted multiple reaction monitoring (MRM) while quadrupole time of flight (Agilent 6530 and 6545) instruments were used for stable isotope tracer analyses. Chromatography was achieved using a 1290 Infinity ultra-high performance LC system (Agilent Technologies, Santa Clara, CA, USA). The mass spectrometers are equipped with a Jet Stream electrospray ionization (ESI) source. For all LC-MS analyses, 5 µL of the sample was injected. Data were analyzed using MassHunter Quant (Agilent Technologies) for steady state analysis and Profinder (Agilent Technologies) for stable isotope tracer analysis. No additional corrections were made for ion suppression. For both LC/MS analyses, the peak intensity (integrated peak area) of each metabolite was identified to calculate its relative abundance compared to the medium only (naïve medium). The values were normalized using the total sum of peak area for all metabolites analyzed for each sample, and differential enrichment of metabolites was calculated for drug and siRNA treatment using fold change and logarithmic analysis. Further statistical analyses were also done using Metaboanalyst online tools.

**Ion pairing method**. Samples were analyzed in negative mode. Multiple reaction monitoring was optimized on authentic metabolite standards. Source gas temperature and flow were set at 150°C and 13 L/min, respectively, nebulizer pressure was set at 45 psi, and capillary voltage was set at 2000V. The autosampler temperature was maintained at 4°C. Metabolite separation was achieved by using a Zorbax Extend C18 column 1.8 µm, 2.1 × 150mm$^2$ with guard column 1.8 µm, 2.1 × 5mm$^2$ (Agilent Technologies) maintained at 35°C. The chromatographic gradient started at 100% mobile phase A (97% water, 3% methanol, 10 mM tributylamine, 15 mM acetic acid, 5 µM medronic acid) for 2.5-min, followed by a 5-min gradient to 20% mobile phase C (methanol, 10 mM tributylamine, 15 mM acetic acid, 5 µM medronic acid), a 5.5-min gradient to 45% C and a 7-min gradient to 99% mobile phase C, which was kept for 4 min. The column was restored by washing with 99% mobile phase D (90% ACN) for 3 min at 0.25 mL min$^{-1}$, followed by an increase of the flow rate to 0.8 mL min$^{-1}$ over 0.5 min and a 3.85-min hold, after which the flow rate was reduced to 0.6 mL min$^{-1}$ over 0.15 min. The column was then re-equilibrated at 100% A over 0.75 min, during which the flow rate was decreased to 0.4 mL min$^{-1}$ and held for 7.65 min. One minute before the next injection, the flow was returned to 0.25 mL/min at 35°C. Retention times and linear detection range were obtained by running authentic standard mixes. Repeat injections of authentic standards were performed throughout the run to observe any retention time shifts and chromatographic quality. The areas under the curve for each sample and metabolite were analyzed to ensure they were below the saturation limit for those metabolites for which range curves were available. Samples were diluted 10-fold and re-run for analytes exceeding the linear range of the instrument.

**Intrada-AA method**. For better coverage of positive ionizing metabolites and amino acids which are not retained in reversed phase, we used a hydrophilic interaction chromatography. Using same parameters for gas temperature and flow, nebulizer pressure and capillary voltage, chromatographic separations of metabolites were achieved using an Intrada Amino Acid column 3 µm, 3.0×150mm (Imtakt Corp, JAPAN). The chromatographic gradient started at 100% mobile phase B (0.3% formic acid in ACN) with a 3 min gradient to 27% mobile phase A (100 mM ammonium formate in 20% ACN / 80% water) followed by a 19.5 min gradient to 100% A at a flow rate of 0.6 ml/min. This was followed by a 5.5 min hold time at 100% mobile phase A and a subsequent re- equilibration time (7 min) before the next injection. The column temperature was maintained at 10°C. Relative concentrations were determined from external calibration curves of standards dissolved in water to ensure that samples analyzed were within the linear range of the instrument. For those that exceeded the linear range 30-fold dilutions were made and samples re-analyzed. For all LC/MS analyses, 5 µL of the sample was injected. Data were analyzed using MassHunter Quant (Agilent Technologies). No additional corrections were made for ion suppression.

**Stable Isotope Tracer analyses (SITA)**. For stable isotope tracer metabolite analysis of glutamine, Hs 578T was maintained as described and treated with DMSO or 1-2 µM UNC for 2 days. DMSO- and UNC-treated Hs 578T cells were then grown in DMEM containing 4 mM $^{13}$C5-labelled glutamine (CML-1822; Cambridge Isotope Laboratories, Inc.) or unlabelled glutamine for 2 h. Samples were collected and extracted as explained above. The same columns, chromatographic methods, and source settings were used for SITA as noted above. Data were collected in high resolution scan mode on both QTOF instruments (6530 and 6545 QTOF). Retention times and linear detection ranges were determined by running authentic standard mixtures. Repeat injections of authentic standards were performed throughout the run to observe any shifts in retention time, interferences or chromatographic quality. The areas under the curve for each sample, metabolite, and isotopes were analyzed and ensured to be below the saturation limit for those metabolites where range curves were available. Stable isotope tracer data were analyzed, and natural abundance matrix correction was performed using Profinder software (Agilent Technologies). Cells were exposed to $^{13}$C-labeled nutrients, and control samples were exposed to natural abundance nutrients to ensure that observed $^{13}$C incorporation into other metabolites is not an interfering ion.
